# Residual cohesin function and PARP1 levels determine PARP inhibitor sensitivity in STAG2-deficient cancers

**DOI:** 10.64898/2026.09.04.749084

**Authors:** Thom M. Molenaar, Zoi Karagiorgou, Sacha Jacobs, Daan L. Tang, Isha Sadal, Martin A. Rooimans, Job de Lange

## Abstract

The STAG1 and STAG2 paralogs are mutually exclusive subunits of cohesin, a large protein complex that controls different aspects of genome organization. Loss-of-function mutations in STAG2 are common in a variety of cancers, and have been associated with increased sensitivity to poly(ADP-ribose) polymerase inhibitors (PARPi). This provides a possible entry point for therapeutic exploitation. However, insights in the underlying mechanism and additional determinants of this sensitivity are currently lacking. Here we show that STAG2-deficient cell lines exhibit variable PARPi sensitivity. While wildtype TP53 enhances PARPi sensitivity in RPE1 cells, STAG2 depletion does not trigger p53 or affect PARPi sensitivity in RPE1-TP53wt cells. PARPi sensitivity is determined by the capacity of STAG1 to compensate for the lack of STAG2. Thus, STAG2 is not uniquely required for the PARPi response, and its loss predominantly sensitizes cells through a general defect in cohesin function. PARP trapping underlies this sensitivity and is strongly influenced by basal PARP1 expression levels and further enhanced by oncogenic signaling. Perturbations in sister chromatid cohesion also enhances PARPi sensitivity, correlating with aggravated cohesion defects and increased chromosome breakage. Mechanistically, cohesin impairment does not appear to compromise repair of nickase-induced DNA lesions, suggesting that cohesin-defective cells are particularly vulnerable to a distinct class of lesions associated with trapped PARP. Together, our findings identify residual cohesin function and PARP1 expression as key determinants of PARPi response and suggest that successful clinical application of PARP inhibitors in STAG2-mutant cancers will require biomarker-guided patient stratification rather than STAG2 mutation status alone.

## Introduction

Cohesin is a ring-shaped protein complex that can hold two DNA strands together. During DNA replication, cohesin establishes sister chromatid cohesion (SCC) to tether the newly formed DNA strands together until their separation in mitosis (Nishiyama 2019; Peters and Nishiyama 2012; Srinivasan et al. 2020). In addition, throughout the cell cycle, cohesin continuously cycles on and off chromatin to establish dynamic DNA contacts between distal elements on the same chromosome (Nishiyama 2019; Rowley and Corces 2018; Yatskevich, Rhodes, and Nasmyth 2019; Perea-Resa et al. 2021; Rittenhouse and Dowen 2024). By controlling DNA contacts in both *trans* and *cis*, cohesin regulates chromosome segregation and genome stability as well as chromatin conformation and gene expression.

In human somatic cells, the cohesin complex consists of a heterodimer of SMC1A and SMC3, the kleisin subunit RAD21, and either STAG1 or STAG2 (Shi et al. 2020). STAG1 and STAG2 are paralogs, associating with cohesin in a mutually exclusive manner (Losada et al. 2000; Sumara et al. 2000). In general, STAG1 and STAG2 control dynamic interactions between cohesin and DNA to promote loop extrusion (Davidson et al. 2019; Hu et al. 2011; Li et al. 2018; Murayama and Uhlmann 2014), and also mediate interactions with other proteins such as the chromatin architecture protein CTCF (Li et al. 2020). Cohesin-STAG1 and -STAG2 largely overlap in their distribution across the genome (Arruda et al. 2020) but also have distinct functions. For example, cohesin-STAG2 is more frequently associated with transient contacts between regulatory DNA elements such as enhancers and promoters, while cohesin-STAG1 more frequently mediates more stable contacts at the boundaries of topologically associating domains (TADs) in concert with CTCF (Alonso-Gil et al. 2023; Wutz et al. 2020; Kojic et al. 2018; Casa et al. 2020; Viny et al. 2019).

Mutations in genes encoding the cohesin subunits and its regulators are common in cancer (Hill, Kim, and Waldman 2016). *STAG2* is the most frequently mutated cohesin gene, with 41% of mutations leading to truncation and 56% causing missense substitutions (Romero-Pérez et al. 2019). *STAG2* mutations occur in multiple cancer types including but not limited to bladder cancer, endometrial cancer, colorectal cancer, and multiple types of leukemia, supporting a tumor-suppressor role for STAG2 (for review see (Waldman 2020; Scott et al. 2025)). STAG2 deficiency has been associated with genome instability by causing DNA replication stress and replication-associated DNA repair defects (Mondal et al. 2019). STAG2 deficiency has also been linked to SCC defects and aneuploidy (Solomon et al. 2011) although this is not consistently observed in patient samples or cells harboring patient-derived mutations in *STAG2* (Balbás-Martínez et al. 2013; Kim et al. 2016). Increasing evidence suggests that STAG2 loss drives tumorigenesis via transcriptional dysregulation, for example by promoting self-renewal over differentiation in leukemia (Viny et al. 2019; Fischer et al. 2024; Smith et al. 2020; Sasca et al. 2019; Mullenders et al. 2015; Galeev et al. 2016), by disrupting differentiation in bladder cancer (Richart et al. 2021) and lung cancer (Ashkin et al. 2025), or by altering gene expression programs linked to metastasis in Ewing sarcoma (Adane et al. 2021; Surdez et al. 2021).

While the exact impact of STAG2 and other cohesin gene mutations on tumorigenesis is still being investigated, vulnerabilities in STAG2 mutant cancer cells have already been identified that might be exploited in novel therapies. Loss of STAG1 is invariably lethal in STAG2-deficient cells (van der Lelij et al. 2017; Bailey et al. 2021) – an example of synthetic lethality due to a complete loss of cohesin function. This makes STAG1 a potential drug target, though developing STAG1-targeting therapies is not straightforward. Another more readily targetable weakness in STAG2-deficient cells is their sensitivity to various types of DNA damage. In multiple cancer cell types, STAG2 loss increases sensitivity to agents that cause DNA replication stress or DNA damage such as topoisomerase inhibitors or poly(ADP-ribose) polymerase (PARP) inhibitors (Tothova et al. 2021; Mondal et al. 2019; Zhou et al. 2023; Bailey et al. 2014; Bailey et al. 2021). In particular PARP inhibition (PARPi) has attracted interest as a potential therapy for STAG2-deficient cancers as it could represent a more selective and less toxic therapy compared to other DNA damaging chemotherapeutics, and several PARP inhibitors are already clinically approved for *BRCA1* mutated breast and ovarian cancer.

The efficacy of PARPi in targeting STAG2-deficient cancer cells has been shown both *in vitro* and in mouse models (Tothova et al. 2021). However, the factors that influence PARPi sensitivity in these cells remain unclear, and linked to this, the exact mechanism of action of PARPi in the context of STAG2 deficiency is not fully understood. Clarifying this will be essential for patient stratification and the clinical success of PARPi in *STAG2* mutated cancers. Here we show that PARPi sensitivity varies widely among different STAG2-deficient cell lines, and depends on PARP1 protein levels and the remaining cohesin functionality imparted by STAG1. Genetic perturbations that affect cohesin, or SCC specifically, lead to PARPi sensitivity in a manner that depends on PARP1 levels. Our results are most consistent with a model in which cohesin promotes the repair of DNA damage caused by PARP trapping, and that factors that exacerbate SCC loss or promote this trapping determine PARPi sensitivity in STAG2-deficient cells.

## Methods

### Cell culture

RPE1-hTERT (hereafter RPE1), J82, UM-UC-3, Colo320, and Caco-2 cell lines were grown in DMEM with 8% FCS, 1mM sodium pyruvate, and penicillin–streptomycin (all from Gibco). RT112, JMSU-1, and K562 cell lines were grown in RPMI with 8% FCS, and penicillin–streptomycin. Cells were maintained at 37°C, 5% CO_2_.

### Gene editing and knockdowns

RPE1 cells with knock-out (KO) of the puromycin resistance gene (left over from the pGRN145 hTERT plasmid), *TP53* KO, and doxycycline (dox) inducible Cas9 (hereafter RPE1 p53KO Tet-Cas9) were described previously (van der Weegen et al. 2021). RPE1 and K562 cells with dox inducible CRISPR interference (CRISPRi) using ZIM3-dCas9 (Alerasool et al. 2020) were made by transducing cells with pTREZ-CRISPRi (for plasmid sequences see Supplementary File 1), selecting cells with 100 µg/mL zeocin (Invivogen), and selecting clones that showed high ZIM3-dCas9 levels by immunoblotting after 72h with dox. Similarly, dox inducible Cas9 was introduced in Colo320, RT112, UM-UC-3, and K562 by transducing cells with pTREZ-Cas9, selecting cells with 100 µg/mL zeocin, and selecting clones that showed high Cas9 levels after 72h with dox. Unless otherwise stated, Cas9 and ZIM3-dCas9 were induced with 500 ng/mL dox. Caco-2, K562, JMSU-1, and J82 cells with constitutive Cas9 were made by transducing cells with lentiCas9-blast (addgene plasmid #52962 (Sanjana, Shalem, and Zhang 2014)) and selecting with 10 µg/mL blasticidin (Invitrogen).

RPE1 p53KO Tet-Cas9 *STAG2* KO cells (Benedict et al. 2020), *ESCO2* mutant cells (van Schie, de Lint, Molenaar, et al. 2023), and *DSCC1* KO cells (van Schie, de Lint, Pai, et al. 2023) were described previously. KO cells were generated either through transfection of synthetic crRNAs or through lentiviral transduction of guide RNAs (gRNAs). An overview of how each KO was generated, including crRNA and gRNA sequences, is provided in Supplementary Table S1. For crRNA transfection in RPE1 p53KO Tet-Cas9 cells, Cas9 expression was first induced with dox for 24h, followed by transfection of 20 nM equimolar crRNA:tracrRNA duplexes (both from IDT) using 1:1000 RNAiMax (Life Technologies). *TP53* was knocked out in RPE1 pTREZ-CRISPRi cells using Cas9 ribonucleotide protein (RNP) transfection. For this, 20 nM equimolar crRNA:tracrRNA duplex with 2 µg Alt-R SpCas9 Nuclease V3 (IDT) was transfected using 1:1000 RNAiMax followed by selection with 10 µM Nutlin-3 (Selleck Chemicals). For knockdowns using CRISPRi, lentiviral vectors expressing two gRNAs simultaneously against the same gene were used as this increases knockdown efficiency (Replogle et al. 2022). Lentiviral gRNA vectors with puromycin resistance were selected using 10 µg/mL puromycin (Sigma-Aldrich) in RPE1 pTREZ-ZIM3-dCas9 cells, 5 µg/mL puromycin in RPE1 p53KO Tet-Cas9 cells, or 1 µg/mL puromycin in K562, RT112, J82, JMSU-1, Colo320, Caco-2, and UMUC3 cells. Lentiviral gRNA vectors with neomycin resistance were selected using 500 µg/mL G418 (Santa Cruz Biotechnology) for RPE1 cells or 800 µg/mL G418 for K562 cells.

For microRNA (miRNA) based knockdowns in Colo320, we used the miR-E system (Fellmann et al. 2013). Lentiviral vectors with miRNAs embedded in the 3’UTR of a dox inducible mCerulean fluorescent protein gene and a blasticidin resistance gene were transduced to Colo320 cells which were selected with 10 µg/mL blasticidin. Clonal cell lines displaying high mCerulean expression by fluorescent microscopy after treatment with 500 ng/mL dox (two clonal cell lines per miRNA and two independent miRNAs per target gene) were selected for further experiments. Sequences of miRNAs can be found in Supplementary Table S1.

### Expression constructs

Plasmid sequences of cDNA expression constructs can be found in Supplementary File 1.

### Lentivirus production

Lentivirus was produced in HEK293T cells using 3^rd^ generation packaging plasmids pRSV-Rev (addgene plasmid #12253), pMD2.G (addgene plasmid #12259), and pMDLg/pRRE (addgene plasmid #12251). Transfer and packaging plasmids were transfected using polyethylemine MW 25k (Polysciences), virus was harvested at 48h and 72h, and filtered through a 0.45 µm syringe filter.

### Drug sensitivity and cell viability assays

Drug details are listed in Supplementary Table S1. For drug sensitivity assays using dox inducible CRISPRi, cells were pre-treated with dox for 48 before seeding in 96 well plates. For all cell lines except K562, 1300 cells/well were seeded in 96 well plates, drug dilutions were added, and cells were incubated for 5 days. For K562, 2600 cells/well were seeded and cells were incubated for 6 days. For experiments involving destabilizing domain (DD) tagged proteins (Banaszynski et al. 2006), 0.5 µM Shield-1 was added simultaneously with drug dilutions. Cell viability was measured using CellTiter-Blue (Promega).

Clonogenic survival assays were performed by seeding 200 cells/well in 6 well plates. Medium and drugs were refreshed every 3-4 days and cells were incubated for 14 days. Cells were stained with crystal violet and colony area coverage was measured using a GelCount imager (Oxford Optronix).

### Immunoblotting

Cells were washed twice with PBS and lysed in SDS lysis buffer (50 mM Tris-HCl pH 6.8, 2% SDS, 10% glycerol) with cOmplete protease inhibitor cocktail (Roche). DNA was sheared by passing lysate through a small gauge needle. Protein concentrations were measured using the DC Protein Assay (Biorad) and lysates were supplemented with 0.1 mM DTT and bromophenol blue before being heated to 95°C for 5 minutes. Proteins were separated on 4–15% Mini-PROTEAN TGX Precast Protein Gels (Biorad) and transferred to 0.45 µm PVDF membrane (Millipore). Membranes were blocked in 5% dry milk in PBS and antibodies (Supplementary Table S1) were incubated overnight in a cold room in 2% dry milk in Tris-buffered saline with 0.1% Tween-20 (TBST). Secondary antibodies coupled to HRP (Supplementary Table S1) were incubated in 2% dry milk in TBST for 1h at room temperature. Membranes were imaged using ECL Prime Western Blotting Detection Reagent (Amersham).

### RT-qPCR and RNA-seq

Total RNA was isolated using the RNeasy Mini kit (Qiagen). For RT-qPCR, 1 µg RNA was converted into cDNA using the iScript cDNA Synthesis Kit (Biorad). qRT-PCR was performed on a LightCycler 480 machine using LightCycler 480 SYBR Green I Master mix (Roche). Primers are listed in Supplementary Table S1. For RNA-seq, library construction and sequencing were performed by GenomeScan (Leiden, The Netherlands) using the NEBNext Ultra II Directional RNA Library Prep Kit. mRNA was enriched from total RNA by oligo(dT) magnetic bead capture, followed by fragmentation, first- and second-strand cDNA synthesis, end repair, adaptor ligation, and PCR amplification. Libraries were sequenced on an Illumina NovaSeq 6000. Both alignment of FASTQ files to the human reference genome (hg38) and gene-level raw read counting were performed using RNA STAR (Dobin et al. 2013). Differential expression analysis was performed using DESeq2 (Love, Huber, and Anders 2014). Differential expression with false discovery rate (FDR) < 0.01 was defined as significant.

### Immunofluorescence

For measuring γH2A.X levels, cells grown on coverslips were washed twice with PBS, fixed for 15 min using 4% formaldehyde in PBS, washed with PBS again, and blocked/permeabilized in PBS with 0.3% Triton X-100 and 3% bovine serum albumin (BSA). Coverslips were washed with PBS, and primary antibody (Supplementary Table S1) was incubated for 2h at room temperature in PBS plus 0.1% Tween-20 and 1% BSA. After three more washes with PBS, Alexa Fluor 488 coupled secondary antibody (Supplementary Table S1) was incubated for 1h at room temperature. Coverslips were mounted in ProLong Gold Antifade Mountant with DAPI (Invitrogen) on slides and imaged using fluorescence microscopy (Leica).

### Sister chromatid cohesion analysis

Cells were treated with 200 ng/mL colcemid for 20 min to arrest cells in metaphase. Cells were subsequently swollen in 0.075 mM KCl, fixed in 3:1 methanol:acetic acid, dropped onto glass slides, and stained in 5% Giemsa solution. For each condition, at least 50 chromosome spreads were blindly assessed for railroad chromosomes and premature chromatid separation.

### Statistical analysis

Statistical analyses were performed using GraphPad Prism (version 10.6.0). The statistical test used for each experiment is indicated in the corresponding figure legend. Differences were considered statistically significant when *P* < 0.05. Statistical significance is denoted as follows: *P* ≥ 0.05 (ns); *P* < 0.05 (*)*; P < 0.01* (**)*; P < 0.001* (***); and *P* < 0.0001 (****).

## Results

### PARPi sensitivity varies among *STAG2* knockout cancer cell lines

We first wanted to determine whether STAG2 loss consistently increases PARPi sensitivity in different cell lines. We therefore knocked out *STAG2* as well as its paralogue *STAG1* in non-transformed human retinal pigment epithelial cells (RPE1-hTERT, hereafter RPE1) lacking p53 (RPE1 p53KO), Colo320 (colorectal cancer, CRC), and K562 (chromic myeloid leukemia, CML) and determined sensitivity to the PARP inhibitors talazoparib and olaparib in independent KO clones. As a control we knocked out *DSCC1* (DNA replication and sister chromatid cohesion 1), a factor important for SCC establishment and error-free DNA replication whose loss causes severe sensitivity to PARPi (van Schie, de Lint, Pai, et al. 2023). STAG2 loss did not sensitize cells to PARPi in either RPE1 p53KO or Colo320 compared to parental or control *OR10A7* (olfactory receptor gene) KO cells (**Figure 1A-B**). Because *STAG2* KO has previously been reported to cause talazoparib sensitivity in RPE1 (Mondal et al. 2019), we repeated our experiment using more long-term colony survival assays (**Supplementary Figure 1**). However, we again did not observe differential sensitivity between parental and *STAG2* KO RPE1 cells using either talazoparib, olaparib, or the topoisomerase I inhibitor camptothecin. We furthermore verified that our RPE1 *STAG2* KO cells indeed lacked functional *STAG2* by showing that *STAG1* is essential in these cells, and that this synthetic lethality could be rescued by re-expressing STAG2 or several STAG2 mutants (**Supplementary Figure 2**).

**Figure 1.**
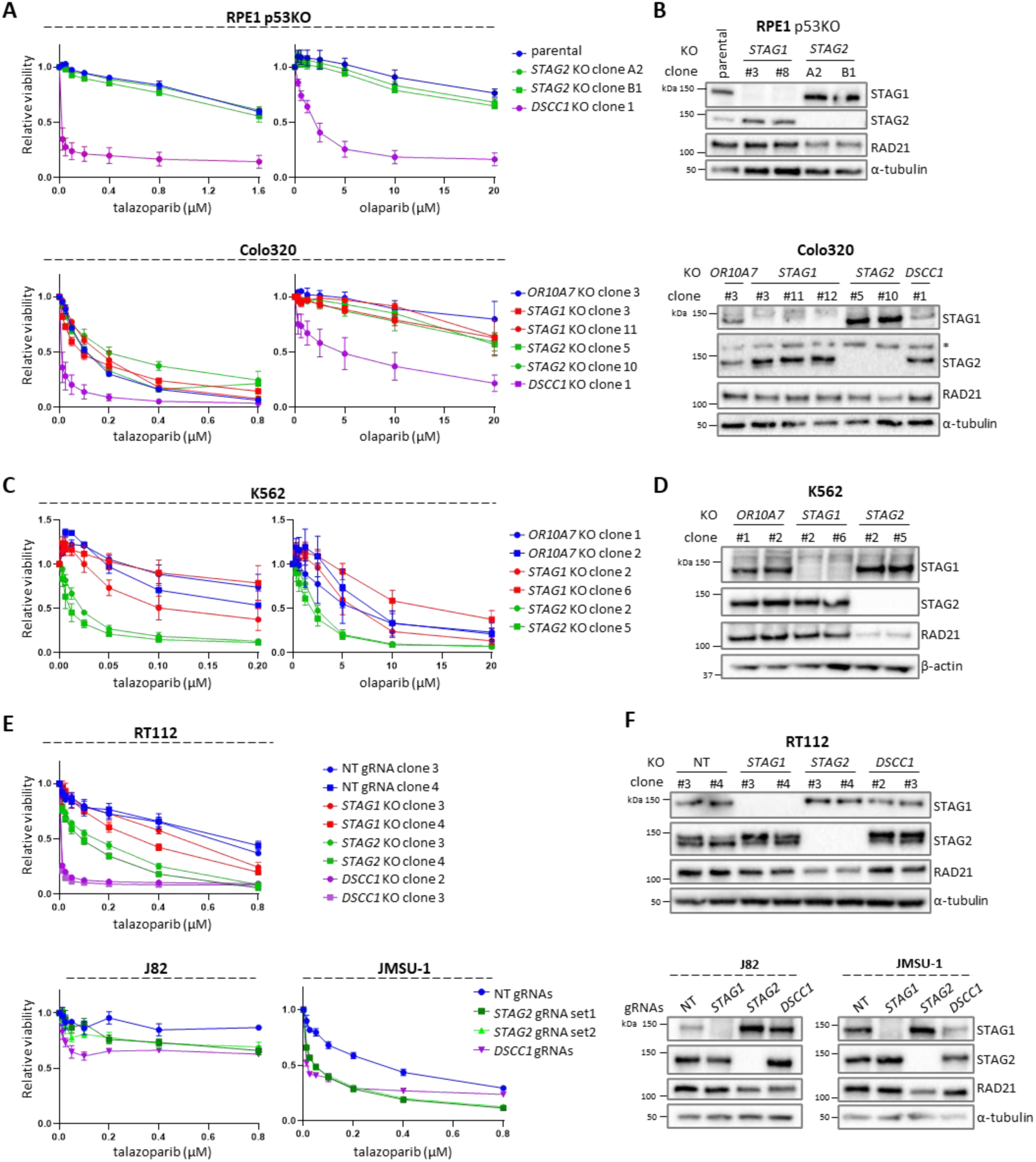
*STAG2* knockout cell lines differ in their sensitivity to PARP inhibitors. **(A, C, D)** PARPi sensitivity assays for *STAG2* KO cell lines. RPE1 p53KO, Colo320, RT112, JMSU-1, and J82 were treated for 5 days with PARPi. K562 was treated for 6 days with PARPi. Error bars represent the standard error of the mean (SEM) of three independent experiments. NT, non-targeting gRNA. *OR10A7* is a non-essential control olfactory receptor gene. **(B, D, E, F)** Whole-cell extract western blots for KO clones, * denotes non-specific bands.

In contrast to RPE1 and Colo320, loss of STAG2 did sensitize K562 cells to PARPi, with talazoparib being more potent than olaparib while STAG1 loss had no effect (**Figure 1C-D**). To exclude any potential clonal biases, we also knocked down *STAG2* in K562 using CRISPR interference (CRISPRi). Again, we observed increased sensitivity after *STAG2* knockdown (KD) although this effect was not as strong as in the *STAG2* KO cells, and also much weaker compared to *BRCA1* KD (**Supplementary Figure 3**). We further observed variable effects of *STAG2* KO on talazoparib sensitivity in urothelial carcinoma cell lines RT112, JMSU-1 and J82 (**Figure 1E-F**).

We also noted that KO of *STAG2* by itself differentially affected viability among cell lines (**Supplementary Figure 4A**). Both K562 and the colon cancer cell line Caco-2 were highly sensitive to acute KO of *STAG2*, but not *STAG1*. Unlike for K562, we were unable to generate *STAG2* KO clones for Caco-2 that could be tested for PARPi sensitivity. K562 *STAG2* KO clones had low levels of RAD21 (see Figure 1D) suggesting that total cohesin levels were decreased. We indeed observed substantial SCC defects in these cells (**Supplementary Figure 4B**) which likely explains the low viability observed in K562 following acute STAG2 loss. Notably, wt K562 cells are characterized by a high STAG2/STAG1 ratio (**Supplementary Figure 4C**). This suggests that certain cell types depend more on STAG2 for maintaining total cohesin levels and cell viability.

Collectively, we conclude that the effect of STAG2 loss on cell viability is highly dependent on the cellular background, and that PARPi sensitivity in *STAG2* KO cells also varies considerably, with some cell lines showing no differential sensitivity compared to wt controls.

### p53 but not STAG2 controls talazoparib sensitivity in RPE1 cells

We next wondered if the resistance of RPE1 p53KO *STAG2* KO cells to talazoparib is influenced by p53 status. For example, it could be hypothesized that STAG2 depletion enhances PARPi sensitivity by eliciting a p53 response, which would be avoided by *TP53* KO. It was previously shown that knocking out *STAG2* itself causes DNA damage and senescence in p53wt RPE1 cells (Mondal et al. 2019). We therefore used a doxycycline (dox)-inducible RNA-targeting RfxCas13d system (Konermann et al. 2018) to efficiently knock down *STAG2* in RPE1 p53wt cells (**Figure 2A**). Because RfxCas13d shows collateral cleavage effects when targeting abundant transcripts (Shi et al. 2023), we used gRNAs targeting the Hygromycin B resistance gene (*HygB*) that is expressed in RPE1 cells as a negative control rather than a non-targeting gRNA. Depending on the gRNA used, *STAG2* KD had minimal or no effect on cell viability while control *RAD21* KD clearly decreased viability (**Figure 2B**). This correlated with a strong up-regulation of p53 protein levels in *RAD21* but not in *STAG2* KD cells (**Figure 2C and D**). In addition, we could readily establish RPE1 p53wt *STAG2* KO clones that grew at a similar rate as wt cells (**Supplementary Figure 5**), confirming that STAG2 loss minimally affects viability in those cells. Importantly, similar as in RPE1 p53KO cells, *STAG2* depletion did not sensitize RPE1 p53wt cells to talazoparib (**Figure 2E**). We also confirmed that depletion of *TP53* using RfxCas13d made RPE1 cells resistant to talazoparib (**Supplementary Figure 6**), highlighting that talazoparib induces a p53 response. Finally, we knocked down *STAG2* and *TP53* simultaneously which again did not sensitize cells to talazoparib (**Figure 2F-H**), recapitulating our initial findings in RPE1 p53KO cells. Taken together, these results show that STAG2 loss does not sensitize RPE1 cells to talazoparib, regardless of p53 status, while p53 loss itself contributes to talazoparib resistance.

**Figure 2.**
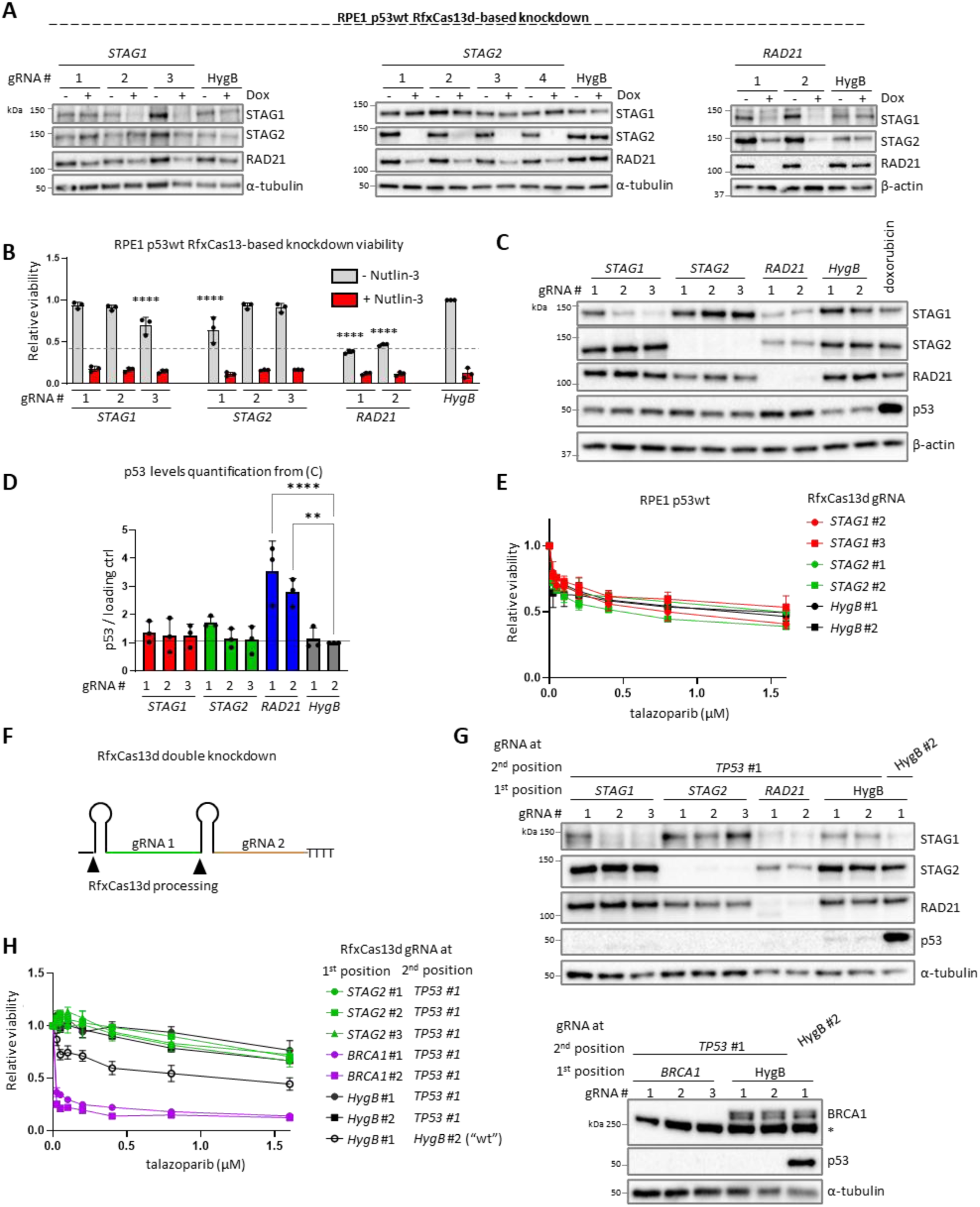
*STAG2* knockdown has no effect on talazoparib sensitivity in p53wt RPE1 cells. **(A)** Whole-cell extract western blots after *STAG1/2* knockdown in RPE1 p53wt cells using dox inducible RfxCas13d (CasRx). A gRNA targeting the hygromycin resistance gene (*HygB*) is used as a control. **(B)** Cell viability 5 days after inducing knockdown with RfxCas13d. Cells were treated with 10 μM Nutlin-3 to test for p53wt status. Comparisons for statistical analysis are against *HygB* -Nutlin-3 control. **(C)** Western blot showing p53 levels after *STAG1/2* and *RAD21* knockdown. Cells treated with doxorubicin (500 nM for 24h) serve as a control for p53 upregulation. **(D)** Quantification of p53 levels by western blotting. **(E)** Talazoparib sensitivity of knockdown cell lines in RPE1 p53wt background. Cells were treated with talazoparib for 5 days. Error bars are SEM of three independent experiments. **(F)** Schematic showing RfxCas13d processing of two gRNAs expressed from a single RNA polymerase III transcript. **(G)** Western blot of RPE1 p53wt cells after double knockdown of *TP53* and either *STAG1/2, RAD21,* or *BRCA1.* **(H)** Talazoparib sensitivity of double knockdown cells. Cells were treated with talazoparib for 5 days. Error bars represent the SEM of three independent experiments. For B and D, error bars represent the SD of three independent experiments. Statistical significance was assessed using one-way ANOVA followed by Dunnett’s multiple comparisons test.

### STAG1 levels in STAG2-deficient cells influence PARPi sensitivity

Having established that PARPi sensitivity varies among different cell lines, we next aimed to uncover which factors determine PARPi sensitivity in STAG2*-*deficient cells. We noted that *STAG2* KO cell lines showed varying degrees of STAG1 protein upregulation (see Figure 1). Increased STAG1 protein levels following the loss or depletion of STAG2 has been observed before in many different contexts (Viny et al. 2019; Wutz et al. 2020; Moronta Gines et al. 2025). In our cell line panel, this apparent compensation by STAG1 was especially prevalent in *STAG2* KO Colo320 cells which were notably also not differentially sensitive to talazoparib compared to their wt counterparts. We verified that STAG1 upregulation is not unique to the monoclonal Colo320 *STAG2* KO lines, as a similar increase in STAG1 levels could be seen in polyclonal Colo320 *STAG2* KO cells (**Supplementary Figure 7A**). *STAG1* mRNA was not upregulated in Colo320 *STAG2* KO cells (**Supplementary Figure 7B**) suggesting that increased mRNA expression or stability does not underly the higher STAG1 levels.

We first wanted to determine if STAG1 protein upregulation following STAG2 loss contributes to talazoparib resistance in Colo320 *STAG2* KO cells. Since a complete loss of STAG1 is lethal in *STAG2* KO cells, we opted for a dox-inducible miRNA knockdown system based on miR-E (Fellmann et al. 2013) to deplete *STAG1* in a titratable manner. We transduced *STAG1* or control miR-E constructs in the polyclonal *STAG2* KO pool and then selected two independent clones that were *STAG2* KO and showed STAG1 depletion following dox treatment (**Figure 3A-B**). Depending on the clone, treating with 25 or 50 ng/mL dox reduced STAG1 expression in *STAG2* KO cells to levels similar as in control Colo320 cells. Importantly, under these STAG1 knockdown conditions, Colo320 *STAG2* KO cells became highly sensitive to talazoparib (**Figure 3C**). Cell viability was mostly unaffected at 25 ng/mL dox, while increasing the dose to 50 ng/mL caused reduced viability, reflecting the synthetic lethal interaction between *STAG1* and *STAG2* (**Figure 3D**). These results indicate that high STAG1 levels contribute to talazoparib resistance in Colo320.

**Figure 3.**
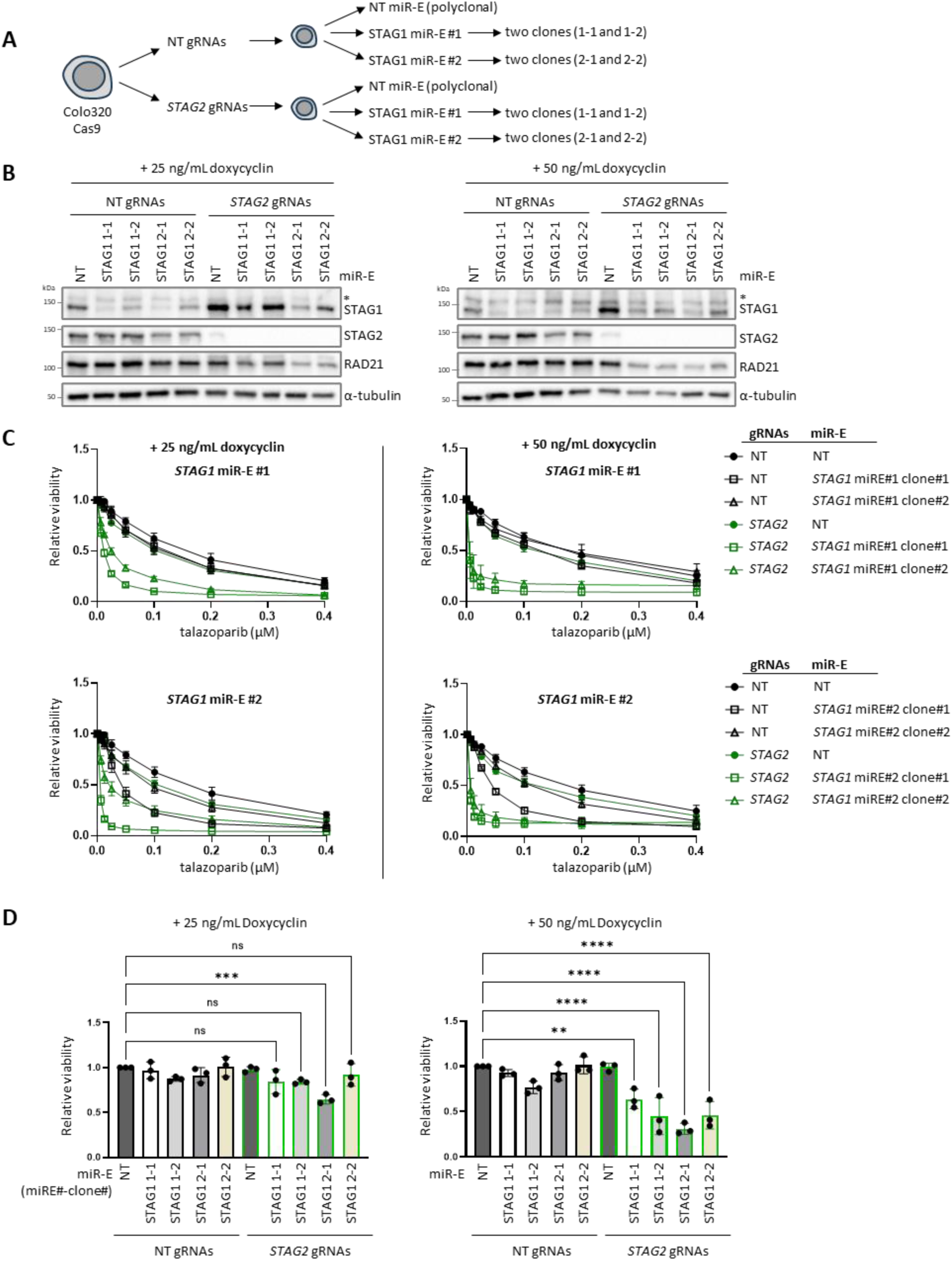
Tuning down STAG1 levels in *STAG2* KO Colo320 cells sensitizes to talazoparib before becoming synthetic lethal with STAG2 loss. **(A)** Overview of how *STAG2* KO Colo320 clones with inducible knockdown of STAG1 were generated. Colo320 cells were transduced with non-targeting (NT) or *STAG2* dual gRNA vectors to make polyclonal KOs, which were then transduced with dox inducible NT or *STAG1* miR-E vectors. Monoclonal cell lines were generated from *STAG1* miR-E transduced cells. **(B)** Whole-cell extract western blot of *STAG2* KO *STAG1* miR-E cells treated with either 25 or 50 ng/mL dox for 72h. Note that there is some STAG2 protein detectable in *STAG2* KO cells with NT miR-E as this is a polyclonal *STAG2* KO. **(C)** Talazoparib sensitivity of *STAG2* KO *STAG1* miR-E cells. Cells were treated with talazoparib for 5 days. Error bars are SEM for three independent experiments. **(D)** Viability of *STAG2* KO *STAG1* miR-E cells. Equal numbers of cells were seeded in a 96-well plate and viability was measured 5 days later. Error bars represent the SD for three independent experiments. Statistical significance was assessed using one-way ANOVA followed by Dunnett’s multiple comparisons test.

We then proceeded to corroborate these findings in talazoparib-resistant RPE1 cells. We created control and *STAG2* KO RPE1 cells with ectopic expression of SMASh-tagged STAG1 as their only source of STAG1 (**Figure 4A**). Using this system, SMASh-STAG1 can be degraded by treating cells with the hepatitis C NS3/4A protease inhibitor telaprevir (TPV) (Chung et al. 2015; Jacobs, Badiee, and Lin 2018). STAG1 is normally expressed as two isoforms through alternative mRNA splicing. We tested both SMASh-STAG1 isoform 1, which is the larger, more abundant isoform, and SMASh-STAG1 isoform 2. We selected two independent clones for each genotype that showed different levels of SMASh-STAG1 in untreated conditions (**Figure 4B**). In absence of TPV, RPE1 *STAG2* KO SMASh-STAG1 cells were already more sensitive to talazoparib (regardless of isoform) compared to both wt and *STAG1* KO RPE1 cells (**Figure 4C**), suggesting that the ectopically expressed SMASh-STAG1 is not as functional as wt STAG1. Importantly, talazoparib sensitivity of the different clones correlated with STAG1 protein levels. Consistent with this, 5 mM TPV treatment made *STAG2* KO SMASh-STAG1 cells highly sensitive to talazoparib. This dose also affected viability in the absence of talazoparib in most clones, again correlating with remaining STAG1 levels (**Figure 4D**). Together we conclude that STAG1 levels in STAG2-deficient cells are an important determinant not only for cell viability but for talazoparib sensitivity as well.

**Figure 4.**
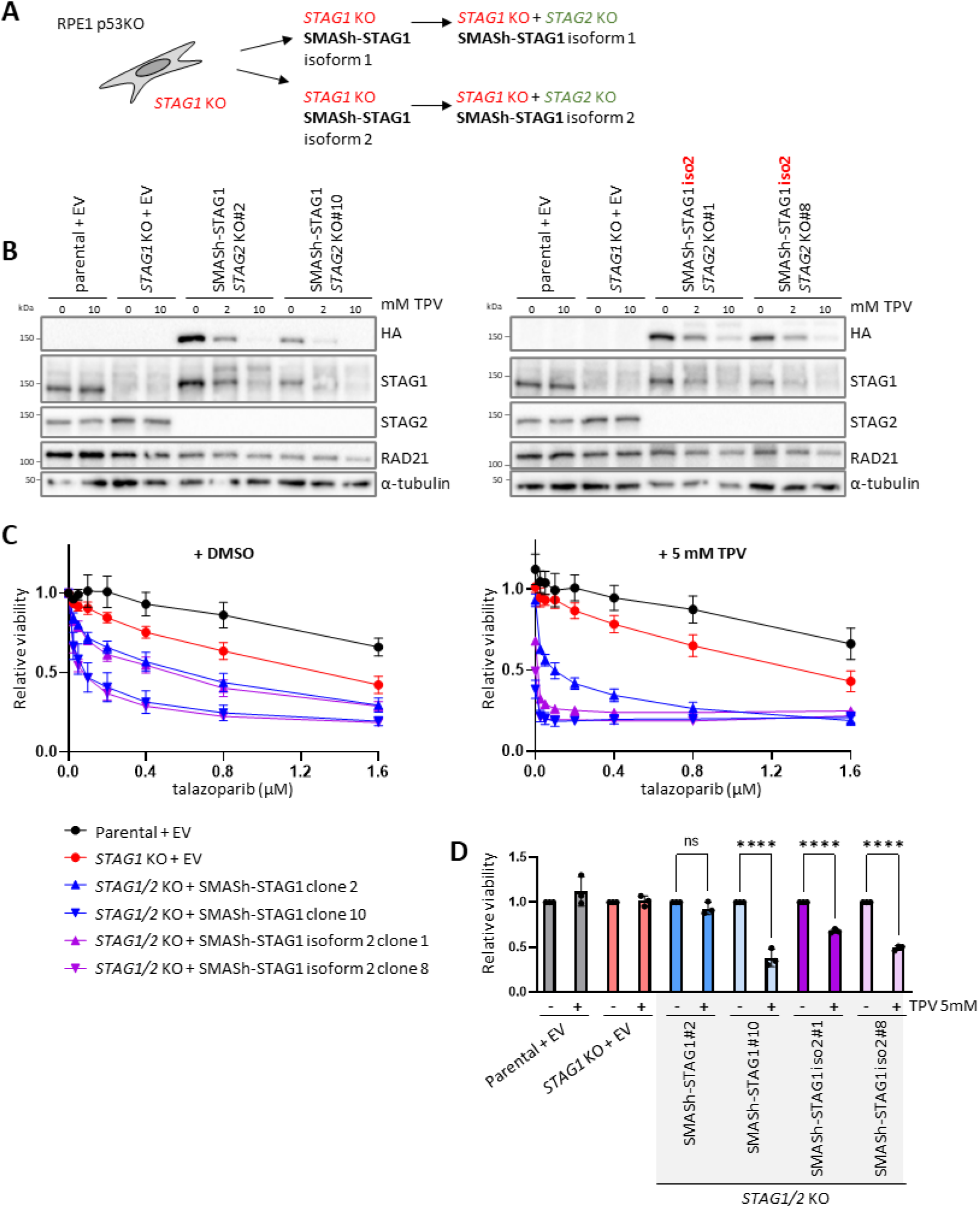
STAG1 depletion sensitizes RPE1 *STAG2* KO cells to talazoparib. **(A)** A monoclonal RPE1 *STAG1* KO cell line was first transduced with degron tagged STAG1 (SMASh-STAG1) either as isoform 1 or 2. Parental and *STAG1* KO cells were also transduced with an empty vector (EV) as control. *STAG2* was subsequently knocked out in SMASh-STAG1 cells and monoclonal cell lines were generated. **(B)** Western blot of SMASh-STAG1 *STAG2* KO cells after treatment with different concentrations of telaprevir (TPV) for 72h. SMASh-STAG1 also contains an HA epitope tag. Note that baseline expression of SMASh-STAG1 varies among clones. **(C)** Talazoparib sensitivity of SMASh-STAG1 *STAG2* KO cells combined with either DMSO or 5 mM TPV. Cell viability was measured after 5 days of treatment. Error bars are SEM for three independent replicates. **(D)** Viability of SMASh-STAG1 *STAG2* KO cells. Equal numbers of cells were seeded in a 96-well plate -/+ 5 mM TPV and viability was measured 5 days later. Error bars are SD for three independent experiments. Statistical significance was assessed using one-way ANOVA followed by Dunnett’s multiple comparisons test.

Our results so far indicate that talazoparib sensitivity in *STAG2* KO cells correlates with remaining cohesin-STAG1 levels. This implies that STAG1 can compensate for STAG2 loss in the context of talazoparib sensitivity. We indeed observed that either ectopic expression of STAG1 or STAG2 could desensitize UM-UC-3 cells to talazoparib (**Supplementary Figure 8**). Although STAG2 may have functions distinct from STAG1 in responding to DNA damage or replication fork stalling (Mondal et al. 2019), our data best support a model in which STAG2-deficient cells become more sensitive to PARPi due to impaired cohesin function when STAG1 cannot sufficiently compensate.

### PARP1 levels influence PARPi sensitivity in STAG2-deficient cells

PARP inhibitors not only inhibit the catalytic activity of PARP but can also cause so-called PARP trapping (Murai et al. 2012). This phenomenon occurs when the PARP enzyme re-binds a DNA damage site without PAR formation (Gopal et al. 2024; Shao et al. 2020), turning the enzyme itself into an impediment for effective DNA repair or replisome progression (Saha et al. 2021; Pommier, O’Connor, and de Bono 2016). Different PARP inhibitors have different abilities to trap PARP, with talazoparib being a highly effective “trapper” (Murai et al. 2014). Consequently, PARP1 depleted cells are resistant to PARPi (Murai et al. 2012; Pettitt et al. 2013).

Following these observations, we reasoned that PARPi resistance may result from low PARP1/2 protein levels in RPE1 cells (Kukolj et al. 2017), as well as in J82 cells (**Supplementary Figure 9**). To test this hypothesis, we expressed PARP1 tagged with a destabilizing domain (DD)(Banaszynski et al. 2006) in RPE1 p53KO cells for Shield-1 inducible PARP1 overexpression (**Figure 5A**). As a control, we overexpressed two PARP1 mutants: PARP1-S568F, which carries a WGR domain mutation that disrupts DNA binding without affecting PARP1 stability (Herzog et al. 2018), and PARP1-E988K which abrogates PARP1’s ability to generate poly-but not mono-(ADP-ribose) (Marsischky, Wilson, and Collier 1995; Rolli et al. 1997). PARP1 overexpression increased talazoparib sensitivity in both wt and *STAG2* KO cells, with a stronger effect in *STAG2* KO cells, resulting in a modest differential sensitivity between the two genotypes (**Figure 5B**). PARP1-E988K further enhanced talazoparib sensitivity in both wt and *STAG2* KO cells, largely eliminating the differential response between them. This could be caused by an inability of PARP1-E988K to generate any PAR at sites of DNA damage, which impairs DNA repair, fork restart and also PARP1 release from DNA, as this is promoted by auto-PARylation (Murai et al. 2012). As expected, PARP1-S568F did not increase talazoparib sensitivity in either background, likely because this mutant cannot be trapped on DNA. These results suggest that low PARP1 expression contributes to talazoparib resistance in STAG2-deficient RPE1 cells.

**Figure 5.**
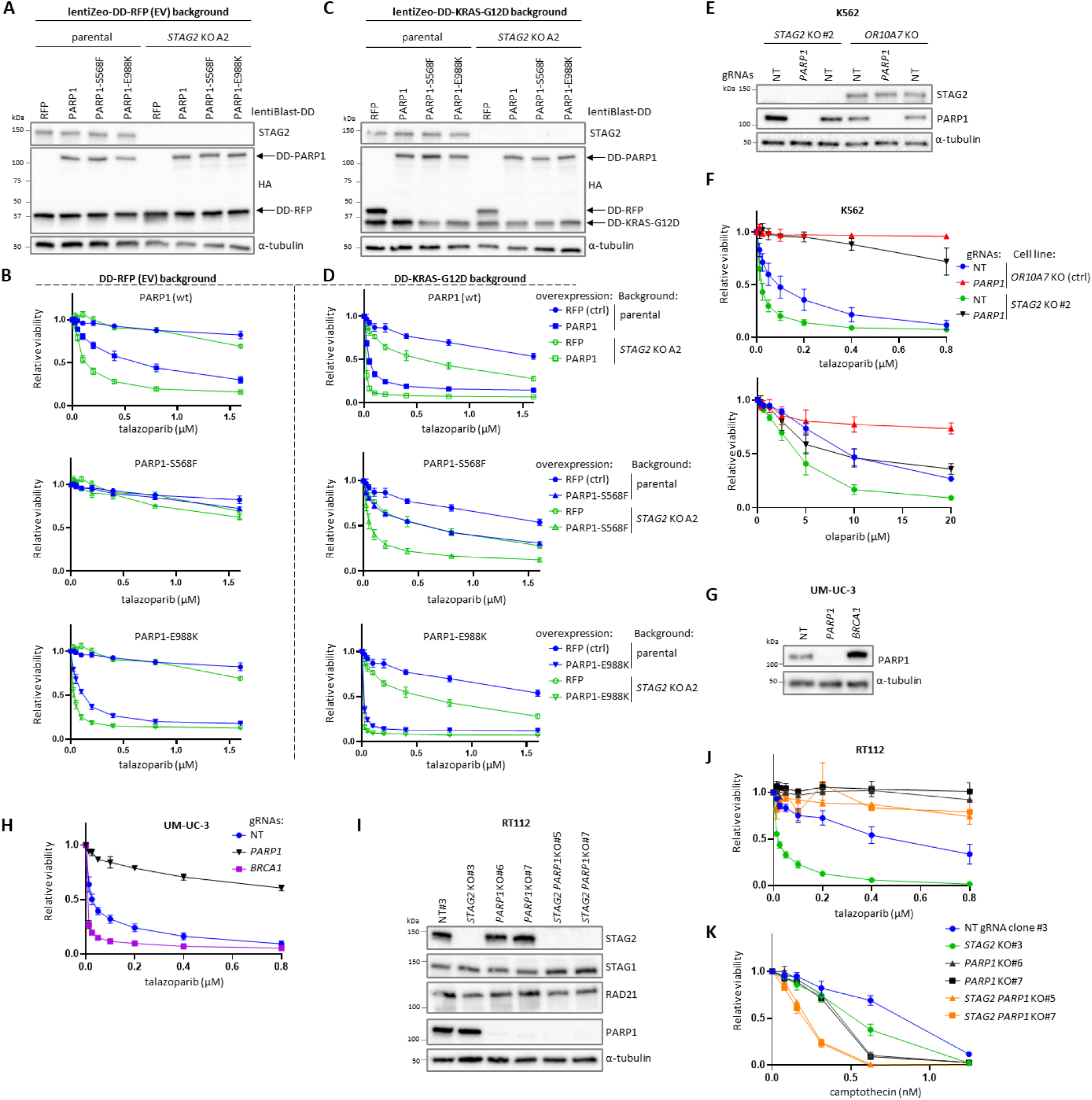
PARP1 levels and KRAS-G12D influence talazoparib sensitivity in *STAG2* KO cells. **(A)** Western blot of RPE1 p53KO *STAG2* KO cells overexpressing Shield-1 inducible DD-PARP1. Cells were transduced with lentiBlast-DD-RFP as a control. Cells were derived from a cell line expressing DD-RFP with a zeocin resistance marker (lentiZeo-DD-RFP) that serves as the empty vector control for KRAS-G12D in C. Cells were treated with 0.5 μM Shield-1 for 72h. **(B)** Talazoparib sensitivity of PARP1 overexpressing RPE1 p53KO *STAG2* KO cells. Cells were treated with 0.5 μM Shield-1 and different concentrations of talazoparib for 5 days. **(C)** Western blot of RPE1 p53KO *STAG2* KO cells with Shield-1 inducible KRAS-G12D and PARP1 overexpression. Cells were treated with 0.5 μM Shield-1 for 72h. **(D)** Talazoparib sensitivity of KRAS-G12D and PARP1 overexpressing RPE1 p53KO *STAG2* KO cells under the same conditions as in B. **(E)** Western blot of *STAG2 PARP1* KO K562 cells. Parental and *STAG2* KO (clone #2) K562 cells with dox inducible Cas9 were transduced with a vector expressing two gRNAs against *PARP1* or two non-targeting (NT) gRNAs resulting in polyclonal *PARP1* KO cell lines. **(F)** PARPi sensitivity of *STAG2 PARP1* KO K562 cells. Cells were treated with PARPi for 6 days. **(G)** Western blot of (polyclonal) *PARP1* KO UM-UC-3 bladder cancer cells with endogenous mutation in *STAG2*. **(H)** Talazoparib sensitivity of *PARP1* KO UM-UC-3 cells. **(I)** Western blot of (monoclonal) *PARP1* and *STAG2* KO RT112 bladder cancer cells. **(J, K)** Talazoparib and camptothecin sensitivity of *PARP1 STAG2* KO RT112 cells. Cells were treated with talazoparib or camptothecin for 5 days. Error bars in B, D, F, H, J, and K represent the SEM for three independent experiments.

In addition to low PARP1 protein levels, RPE1 cells are further characterized by minimal oncogenic signaling due to their untransformed nature, distinguishing them from most cancer cell lines. To assess whether oncogenic signaling contributes to talazoparib sensitivity, we combined PARP1 overexpression with ectopic Shield-1 inducible expression of mutant hyperactive KRAS-G12D. Induction of KRAS-G12D alone led to a modest differential talazoparib sensitivity in *STAG2* KOs compared to parental RPE1 (**Supplementary Figure 10**). Notably, combined overexpression of KRAS-G12D and PARP1 produced a markedly enhanced sensitivity to talazoparib compared to either perturbation alone, with STAG2-deficient cells exhibiting extreme hypersensitivity (**Figure 5C–D**). Cells expressing PARP1-S568F also displayed increased talazoparib sensitivity, which may reflect residual trapping activity or interference with endogenous PARP1 function. These data indicate that both high PARP1 expression and strong oncogenic signaling lead to talazoparib sensitivity in RPE1 cells, with these effects being more pronounced in the context of STAG2 deficiency.

We next assessed the contribution of PARP1 levels to talazoparib sensitivity in additional STAG2-deficient cell line models. In wt K562, we first verified that loss of either *PARP1* alone or *PARP1* and *PARP2* together confers resistance to talazoparib (**Supplementary Figure 11A-C**) similar as has been reported before (Murai et al. 2012; Pettitt et al. 2013). In the context of STAG2 deficiency, *PARP1* loss in K562 also led to resistance to both talazoparib and olaparib (**Figure 5E-F**). Furthermore, *PARP1* KO conferred talazoparib resistance in the urothelial carcinoma cell line UM-UC-3 which has an endogenous truncating mutation in *STAG2*, as well as in RT112 *STAG2* KO cells (**Figure 5G-J**). This effect was specific to PARPi as *PARP1/STAG2* double KO cells were highly sensitive to camptothecin (**Figure 5K**). Notably, *PARP1* KO did not strongly affect UM-UC-3 viability (**Supplementary Figure 11D**). Similarly, loss of *BRCA1*, but not of *STAG2*, decreased viability in RPE1 p53KO *PARP1* KO cells (**Supplementary Figure 11E**). This indicates that STAG2 loss differs from BRCA1 loss in that it does not create a synthetic lethal dependency on PARP1. Collectively, these data indicate that PARP1 levels are a major determinant for talazoparib sensitivity in STAG2-deficient cells.

### Defective SCC causes talazoparib sensitivity

We next sought to determine how cohesin perturbation drives talazoparib sensitivity. PARPi classically targets cells with defects in homologous recombination (HR) where the sister chromatid acts as a template for repair. Cohesin has a multi-facetted role in HR which includes keeping sister-chromatids in proximity through SCC as well as roles in checkpoint signaling and chromatin organization through loop extrusion around DNA double-stranded breaks (DSBs) (reviewed in (Hou et al. 2022; Litwin, Pilarczyk, and Wysocki 2018)). Both non-cohesive cohesin and SCC ultimately contribute to HR by guiding homology search towards the sister chromatid (Teloni et al. 2025). Given the well-established role of cohesin in HR, we first asked whether cohesin disruption via routes other than STAG1 and STAG2 depletion sensitizes cells to talazoparib. To this end, we depleted the cohesin loader MAU2 (Ciosk et al. 2000; Murayama and Uhlmann 2014) in RPE1 p53KO cells using dox-inducible CRISPRi (**Figure 6A**). MAU2 KD sensitized RPE1 cells to talazoparib and this effect was enhanced when PARP1 was overexpressed (**Figure 6B**). MAU2 KD cells were also sensitive to olaparib but not to veliparib, a PARP inhibitor that does not cause trapping (**Figure 6C**), again demonstrating that PARPi sensitivity of cohesin-defective cells results from PARP trapping. MAU2 KD cells were also sensitive to camptothecin (**Figure 6C**), and to a lesser extent to the DNA interstrand crosslinkers mitomycin C and cisplatin (**Supplementary Figure 12**). These results are consistent with previous findings that heterozygous inactivation of *SMC3* or *RAD21* sensitizes cells to talazoparib (Tothova et al. 2021) and that cohesin perturbation in general sensitizes cells to DNA damage.

**Figure 6.**
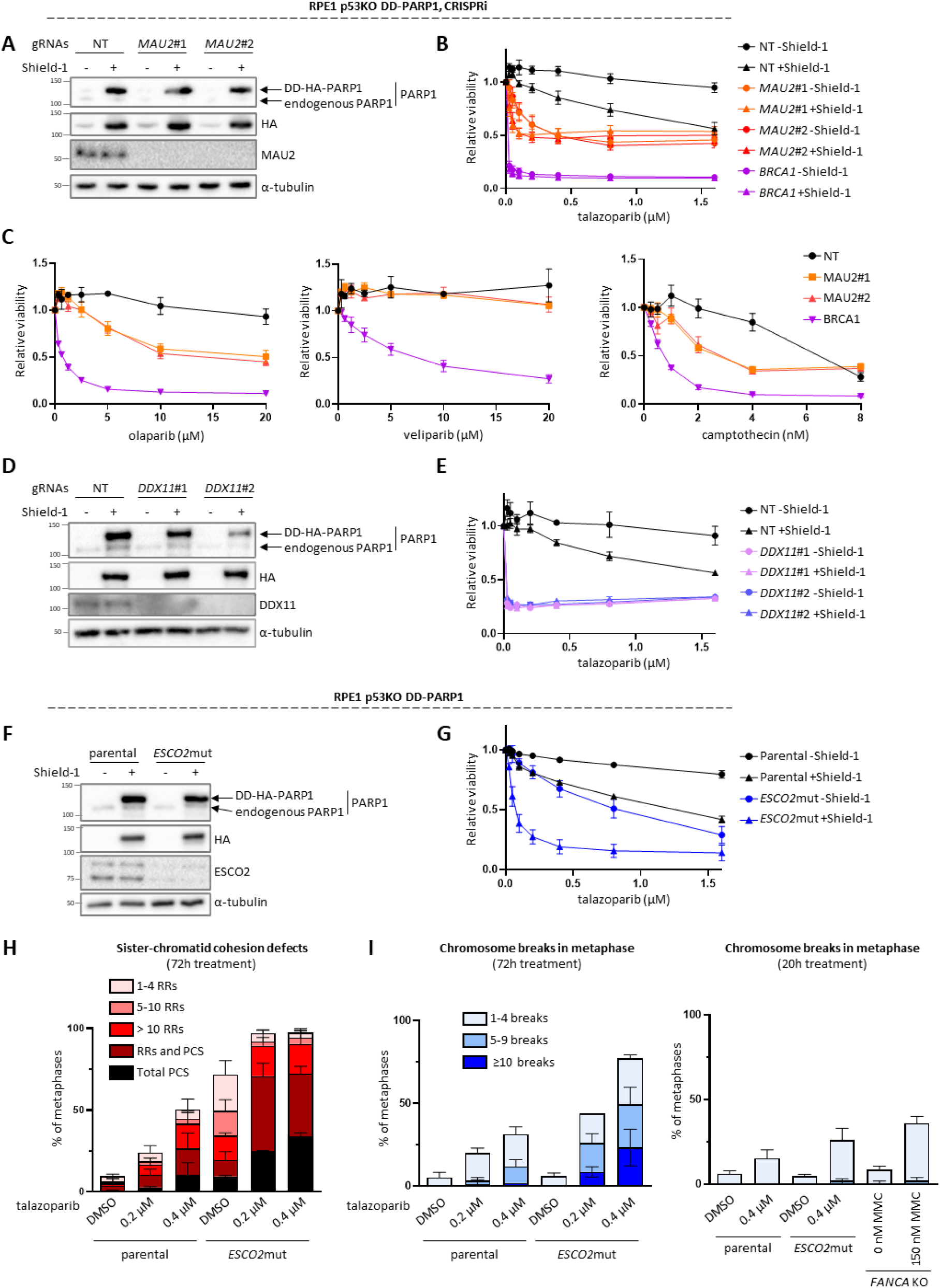
Perturbing cohesin and SCC factors other than STAG1/2 sensitizes to talazoparib. **(A)** Western blot of RPE1 cells with dox inducible CRISPRi against *MAU2* and Shield-1 inducible overexpression of HA-tagged PARP1. Cells were treated with dox and Shield-1 (0.5 μM) for 72h. **(B, C)** Drug sensitivity of *MAU2* KD cells. **(D)** Western blot of RPE1 cells with CRISPRi against *DDX11*. **(E)** Talazoparib sensitivity of *DDX11* KD cells. **(F)** Western blot of RPE1 cells with hypomorphic mutation in *ESCO2* (*ESCO2* mut) and Shield-1 inducible PARP1 overexpression. **(G)** Talazoparib sensitivity of *ESCO2* mut cells. **(H)** Sister chromatid cohesion analysis of PARP1 overexpressing *ESCO2*mut cells treated with talazoparib. RR are railroad chromosomes. PCS, premature chromatid separation. **(I)** Quantification of metaphase spreads containing broken chromosomes. Cells in H and I were pre-treated for 24h with Shield-1 and then treated with talazoparib for 20h or 72h as indicated. Error bars represent SD for three independent experiments. At least 50 metaphases were scored in each condition. For B, C, E, and G, cells were treated with drugs for 5 days. Error bars are SEM for three experiments.

Next we asked whether talazoparib sensitivity arises from perturbation of non-cohesive cohesin (i.e., cohesin engaged in loop extrusion and *cis* DNA contacts) or from disruption of SCC (mediated by cohesive cohesin). SCC can be selectively disrupted by depleting dedicated cohesion establishment factors, such as DEAD/H-Box Helicase 11 (*DDX11*) or Establishment of Sister Chromatid Cohesion N-acetyltransferase 2 (*ESCO2*) (van Schie, de Lint, Molenaar, et al. 2023; van Schie et al. 2020). CRISPRi-mediated *DDX11* KD in RPE1 p53KO cells severely sensitized cells to talazoparib even without PARP1 overexpression (**Figure 6D-E**). Likewise, RPE1 p53KO cells harboring a mutation in *ESCO2* (*ESCO2*mut) that causes SCC defects (van Schie, de Lint, Molenaar, et al. 2023) were also sensitive to talazoparib, particularly in combination with PARP1 overexpression (**Figure 6F-G**). Talazoparib has previously been reported to cause premature SCC loss in cancer cells (Kukolj et al. 2017). In line with this, we observed that talazoparib treatment for 72h induced SCC defects in RPE1 p53KO cells overexpressing PARP1, which was exacerbated in *ESCO2*mut cells (**Figure 6H**). This coincided with an increased number of metaphases with chromosome breaks (**Figure 6I**). These breaks likely reflect the persistence of unresolved DNA lesions into mitosis. Notably, talazoparib treatment for 20h, in which cells can only have experienced a single S-phase, resulted in much fewer breaks in both wt and *ESCO2*mut cells (**Figure 6I**) which supports a model where consecutive S-phases in the presence of talazoparib drive fork collapse and DSB formation (Simoneau, Xiong, and Zou 2021). Thus, impairments of cohesive cohesin result in increased talazoparib sensitivity, which correlates with further aggravated SCC defects and increased chromosome breakage.

Finally, we sought to clarify why cells rely on cohesin/SCC to deal with PARPi. We considered several mechanisms that are not mutually exclusive. In addition to its direct role in SCC and DNA repair, cohesin can also potentially affect DNA damage responses more indirectly by affecting the expression of genes involved in DNA repair. RNA-seq in RPE1 p53KO cells revealed that MAU2 depletion affected several hundred genes (**Supplementary Figure 13A-C**). No gene ontology (GO) terms related to DNA repair were enriched in the differentially regulated genes, and only four down-regulated genes had a GO term related to DNA repair (*ERCC6*, *FAP100*, *LIG3*, and *RECQL*). This makes it unlikely that the talazoparib sensitivity of MAU2 KD cells is primarily driven by indirect transcriptional effects on DNA repair genes. A transcriptional mechanism has also been proposed for DNA damage sensitivity in STAG2-deficient cells, whereby STAG2 loss leads to increased *KMT5A* expression and elevated H4K20me1 levels which impairs BRCA1–BARD1 recruitment (Zhou et al. 2023). Because RT112 *STAG2* KO, but not RPE1 *STAG2* KO cells, are sensitive to talazoparib, we examined *KMT5A* RNA levels in RT112 cells. However, *KMT5A* RNA levels were not increased in RT112 *STAG2* KO cells (**Supplementary Figure 13D**), arguing against involvement of this pathway in our model.

Our results so far indicate that talazoparib mainly affects cohesin-defective cells through PARP trapping. Trapped PARP has been reported to act as a direct roadblock for replication which can be removed by specialized proteases such as SPRTN (Saha et al. 2021). Accordingly, trapped PARP has been reported to cause fork stalling in the absence of SPRTN (Saha et al. 2021). Cohesin may be particularly needed to deal with those impediments, as cohesin accumulates at stalled replication forks where it promotes fork stability, protection, and restart (Morales et al. 2020; Tittel-Elmer et al. 2012; Delamarre et al. 2020; Frattini et al. 2017). We therefore wanted to determine if SPRTN depletion exacerbates talazoparib sensitivity in cohesin-defective cells. Using CRISPRi in RPE1 p53KO cells, we found that *SPRTN* KD did not further enhance talazoparib sensitivity in MAU2 depleted cells (**Supplementary Figure 14**). Together with recent studies that have highlighted that PARP trapping does not involve a prolonged physical interaction between PARP and DNA (Gopal et al. 2024), this could suggest that trapped PARP does not form a physical roadblock to the replication fork. Alternatively, there could be other proteases that remove trapped PARP in RPE1 in absence of SPRTN.

Besides direct effects on fork stability, PARPi causes genome instability by preventing or delaying single-stranded break repair (SSBR), causing replication fork collapse and DSBs in the following S-phase (Simoneau, Xiong, and Zou 2021; Bryant et al. 2005; Farmer et al. 2005). In addition to ssDNA breaks, PARPi also causes post-replicative ssDNA gaps which critically require BRCA1/2 for repair (Cong et al. 2021). The source of these ssDNA gaps is likely PRIMPOL-mediated replication repriming (Tirman et al. 2021) or defective lagging strand/Okazaki fragment maturation (Vaitsiankova et al. 2022). Both DSBs and post-replicative ssDNA gaps can be repaired through mechanisms involving SCC, namely HR for DSBs and template switching for ssDNA gaps, which in budding yeast has been show to depend on SCC (Tittel-Elmer et al. 2012). To determine if cohesin is required for gap repair, we introduced ssDNA gaps in RPE1 p53KO *MAU2* KD cells using either DNA alkylating agents or Shield-1 inducible APOBEC3A overexpression, which both induce gaps by PRIMPOL-dependent repriming (Kawale et al. 2024; Jahjah et al. 2024; Fingerman et al. 2024). In addition, we treated cells with a FEN1 inhibitor (FEN1-IN-1) to impair Okazaki fragment maturation. *MAU2* KD cells showed little or no differential sensitivity to these gap-inducing conditions compared to control cells (**Figure 7A-C**) indicating that cohesin is largely dispensable for post-replicative ssDNA gap repair in RPE1 p53KO cells. We then wanted to determine if persistent DNA nicks and consequent fork collapse in S-phase drive PARPi sensitivity in *MAU2* KD cells. To test this, we expressed grazoprevir (GZV) inducible nickases in RPE1 p53KO cells, either based on a nickase mutant of *Staphylococcus aureus* Cas9 (nSaCas9) (**Figure 7D-E**) or on the monomeric nickase restriction enzyme Nb.Mva1269I (**Supplementary Figure 15A**). We reasoned that inducing DNA nicks in this way could overload single-strand break repair pathways and cause fork collapse. Both systems induced prominent γH2A.X in a proportion of cells indicating that nickase expression led to a DNA damage response involving ATM/ATR (**Figure 7F and Supplementary Figure 15B**). *BRCA1* KD cells were sensitive to both nickase systems, consistent with the model that nickase-induced DNA lesions promote replication fork collapse and require HR for repair (**Figure 7G and Supplementary Figure 15C**). However, *MAU2* KD cells were not or only mildly more sensitive compared to control cells depending on the nSaCas9 gRNA. These results indicate that cohesin is not required for ssDNA repair and that DNA nickases do not recapitulate the effects of talazoparib on MAU2 depleted cells, suggesting that impaired SSBR does not underlie PARPi sensitivity of cohesin-deficient cells.

**Figure 7.**
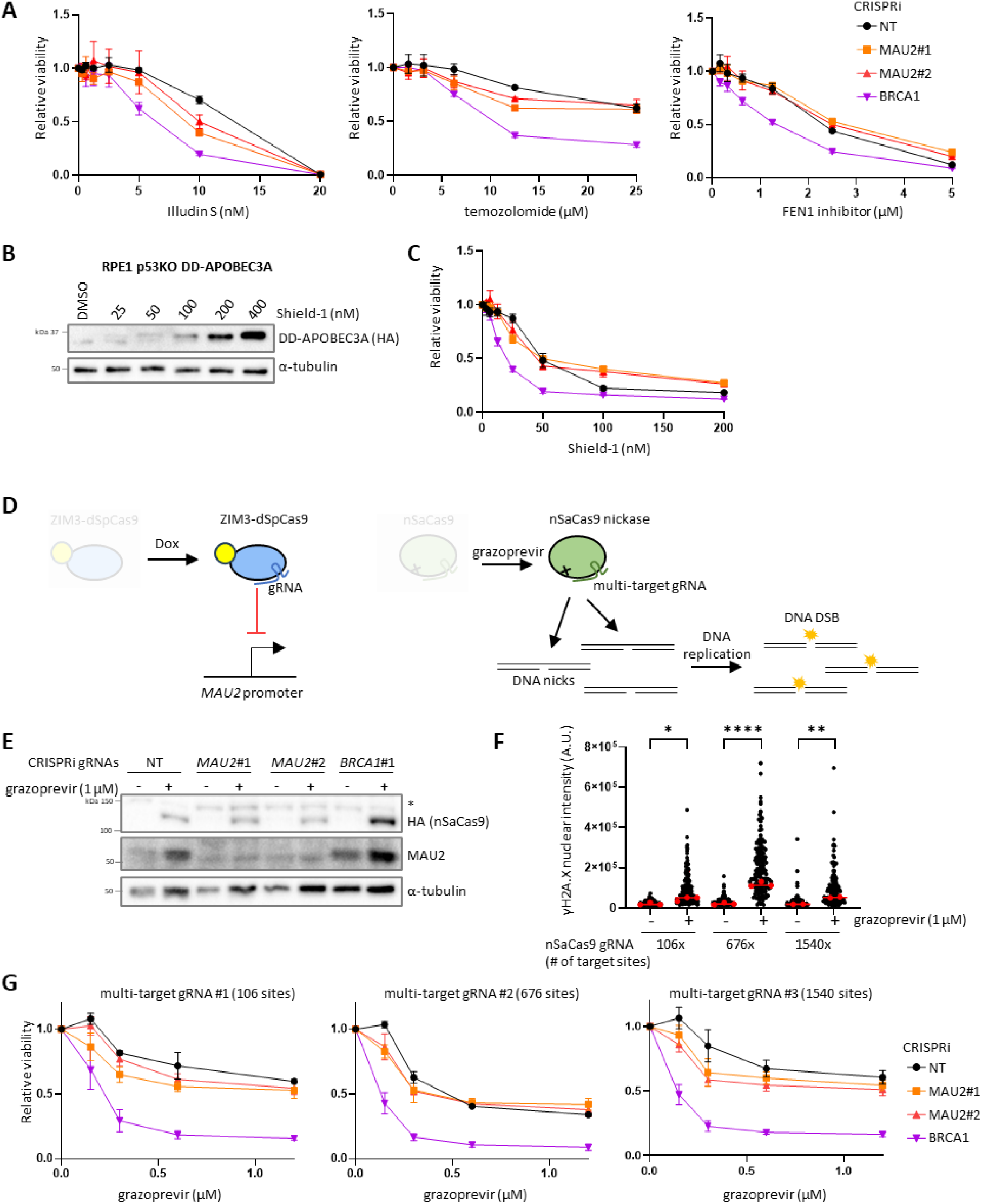
Cohesin depletion does not sensitize cells to post-replicative ssDNA gaps or nickase expression. **(A)** Sensitivity of RPE1 *MAU2* KD cells to DNA alkylating agents temozolomide and illudin S, and FEN1 inhibitor (FEN1-IN-1). **(B)** Western blot of RPE1 cells expressing DD-APOBEC3A treated with different concentrations of Shield-1 for 48h. **(C)** Shield-1 sensitivity of RPE1 DD-APOBEC3A *MAU2* KD cells. **(D)** Overview of RPE1 cells with dox inducible ZIM3-dSpCas9 (CRISPRi) and GZV inducible nSaCas9 (nickase) systems. SpCas9 gRNAs are used for knockdowns, SaCas9 gRNAs targeting multiple times in the genome to generate DNA nicks. **(E)** Western blot of nSaCas9 induction in CRISPRi cells. Cells were treated with GZV for 72h. **(F)** Quantification of γH2A.X immunofluorescence after nSaCas9 expression in cells with different multi-target gRNAs. Cells were treated with GZV for 72 h. Each red dot represents the median γH2A.X intensity of at least 100 nuclei from one independent replicate, and the red bar represents the median of the independent replicates. Statistical significance was assessed by one-way ANOVA followed by Dunnett’s multiple comparisons test, using the median value from each independent replicate as the statistical unit. **(G)** GZV sensitivity of nSaCas9/CRISPRi cells. For A, C, and G, cells were treated with drugs for 5 days. Error bars are SEM for three experiments.

## Discussion

PARP inhibitors are currently clinically approved for use in breast, ovarian, prostate, and pancreatic cancers with defects in HR (Zeng et al. 2024; Hage Chehade et al. 2025). Extending their use to any *STAG2* mutant cancer type has the potential for a significant clinical benefit. Sensitivity of STAG2-deficient cancer cells to in particular talazoparib has been observed in vitro and in xenograft models. Here we extended the panel of STAG2-deficient cell line models, determined factors that control PARPi sensitivity, and revealed the molecular mechanisms behind PARPi sensitivity in STAG2-deficient cells.

We find that STAG2-deficient cell lines strongly differ in their response to PARPi. An important determinant for PARPi sensitivity is the remaining cohesin functionality after STAG2 loss. This is consistent with the observation that perturbation of cohesin function through other means, such as heterozygous *SMC3* or *RAD21* inactivation (Tothova et al. 2021) or *MAU2* depletion (Figure 6), also increases PARPi sensitivity. In contrast, STAG2 has been proposed to play a distinct role from STAG1 in responding to DNA damage and replication stress (Mondal et al., 2019). In our hands, RPE1 STAG2 KO cells displayed moderate talazoparib sensitivity, but only In the context of PARP1 overexpression (Figure 3B). While this might be explained by a specialized role for STAG2 in coping with DNA damage induced by PARP trapping, it seems more likely that excessive PARP trapping increases cellular dependence on cohesin in general, thereby revealing even subtle defects caused by STAG2 loss.

Importantly, our results indicate that STAG1 can largely compensate for STAG2 loss in determining the PARPi response. In some cell lines (such as Colo320) STAG1 is strongly upregulated upon STAG2 loss, while in others (such as K562) the increase is minimal. The mechanism underlying this differential STAG1 upregulation has yet to be determined. We showed that partial depletion of STAG1 in *STAG2* KO Colo320 sensitized cells to talazoparib, confirming that elevated STAG1 levels contribute to PARPi resistance. Furthermore, complementation with either STAG1 or STAG2 enhanced talazoparib resistance in UM-UC-3 cells, which harbor a natural mutation in *STAG2.* Together, these findings indicate that both STAG1 and STAG2 contribute to PARPi resistance. Thus, while we cannot entirely exclude the possibility that STAG2-cohesin is somewhat more effective than STAG1-cohesin in this role, our results point to a general reduction in cohesin function, particularly when SCC is compromised, as the primary determinant of PARPi sensitivity, rather than a unique requirement for STAG2.

Our results indicate that PARP trapping rather than the loss of PARP activity underlies the sensitivity of STAG2- or cohesin-defective cells to PARPi. By contrast, BRCA1-deficient cells were sensitive to both genetic loss of PARP1 and to veliparib, a PARP inhibitor that does not induce PARP trapping (Supplementary Figure 11E and Figure 6C, respectively). This has important implications for any future clinical use of PARPi in *STAG2* or cohesin mutant cancers, as it suggests that weak or medium PARP “trappers” such as veliparib and olaparib will have limited efficacy. Furthermore, PARP levels are a key determinant for PARPi sensitivity in STAG2-deficient cells (Figure 5, Supplementary Figure 11) and should be considered when designing therapies. Finally, loss of PARP1 expression through e.g. mutations or epigenetic silencing could be a potential therapy resistance mechanism in *STAG2* mutant cancer cells. In this case, treatment with other DNA damaging agents such as camptothecin (Figure 5K) could be considered.

How does PARP trapping mechanistically target cohesin- or SCC-defective cells? Trapped PARP-DNA complexes have been proposed to impede replication forks, requiring SPRTN or other proteases for removal (Saha et al. 2021). It is however not entirely clear if trapped PARP indeed acts as a roadblock for replication. DNA fiber analyses with talazoparib show mostly unchanged or even increased fork progression (Vaitsiankova et al. 2022; Pai Bellare et al. 2025; Frederick et al. 2025). Furthermore, PARP trapping does not reflect continuous DNA binding but rather an increased likelihood of repeated DNA rebinding in the absence of auto-PARylation, which normally promotes PARP dissociation (Gopal et al. 2024; Shao et al. 2020). We found that SPRTN depletion did not affect talazoparib sensitivity in *MAU2* KD cells. This argues against a model in which cohesin is needed to deal with stalled forks that result from trapped PARP. This may indicate that trapped PARP does not efficiently stall forks, that alternative proteases remove trapped PARP in RPE1 cells, or that cohesin is largely dispensable for fork recovery under these conditions.

HR-defective cells are highly sensitive to PARPi which lies at the basis of the clinical use of PARPi in HR-defective cancers. Cohesin has a well-established role in HR by keeping sister-chromatids together and by chromatin organization around DSBs to guide homology search (Teloni et al. 2025; Hou et al. 2022; Litwin, Pilarczyk, and Wysocki 2018). Cohesin furthermore facilitates the recruitment of BRCA2 and RAD51 through the cohesin associated factors PDS5A and PDS5B (Morales et al. 2020; Couturier et al. 2016; Brough et al. 2012). The dependence on intact HR in PARPi treated cells has been linked to impaired SSBR or more recently to the formation of ssDNA gaps behind replication forks, increasing the chance for collapse in the subsequent S-phase (Simoneau, Xiong, and Zou 2021; Bryant et al. 2005; Farmer et al. 2005; Cong et al. 2021; Tirman et al. 2021; Vaitsiankova et al. 2022). A simple model is therefore that cohesin perturbed cells are sensitive to PARPi because cohesin promotes HR-mediated repair of collapsed replication forks (nick or ssDNA gap in leading strand) or DSBs behind the fork (nick or ssDNA gap in lagging strand). We tested this model by determining if MAU2 depleted cells are sensitive to the expression of DNA nickases but surprisingly there was only a minor effect. Although more work is required to determine what exactly cohesin’s role is in dealing with DNA lesions induced by these nickases, this suggests that the effects of talazoparib or PARP trapping are not solely explained by the generation of replication-associated DSBs. For example, talazoparib induced SCC weakening may further impair both HR and faithful chromatid separation, and contribute to viability loss in MAU2 depleted cells.

Another possibility is that replication-associated DSBs caused by PARP trapping are repaired differently from nickase induced DSBs. In this regard, it is noteworthy that MAU2-depleted cells were also considerably more sensitive to camptothecin than to the DNA interstrand cross-linkers MMC and cisplatin. Although differences in the kinetics and complexity of the lesions induced by these agents are also likely to contribute to the observed phenotypes, these findings raise the possibility that DSBs associated with covalently trapped TOP1 or dynamically “trapped” PARP have a greater dependency on cohesin for their repair than DSBs generated by DNA nickases or interstrand cross-links. While HR is involved in the repair of all these lesions, as illustrated by the sensitivity to BRCA1 depletion in our experiments, cohesin may play a more prominent role when replication forks encounter bulky (but dynamic in the case of PARP1) nucleoprotein complexes. Future studies will be required to establish whether cohesin preferentially contributes to the HR-mediated repair of bulky nucleoprotein DNA lesions and to define the molecular mechanisms underlying this potential specificity.

In summary, our work points to a model where a reduction in cohesin and SCC levels underlies PARPi sensitivity in STAG2-deficient cells. PARPi sensitivity in STAG2-deficient or otherwise cohesin-depleted cells depends on PARP trapping rather than the loss of PARP catalytic activity. A low dependence on STAG2 for maintaining total cohesin levels, upregulation of STAG1 levels, or reduced PARP1 expression can each contribute to talazoparib resistance following STAG2 loss. The response to PARPi is therefore more varied in STAG2-deficient cancer cells compared to those lacking *BRCA1*. It will therefore be important to determine if specific cancer types with frequent mutations in *STAG2* or other cohesin subunits are more consistently sensitive to talazoparib, or if successful clinical implementation of PARPi for *STAG2* mutant cancers requires stratification of individual patients using biomarkers for functional cohesin or PARP1 expression levels.

## Supporting information

Supplementary File 1 - plasmid maps

## Acknowledgements

We thank Linda Smit for sharing K562 cells and Michiel van der Heijden for sharing bladder cancer cell lines RT112, UM-UC-3, J82 and JMSU-1.

## References

Adane, Biniam, Gabriela Alexe, Bo Kyung A. Seong, Diana Lu, Elizabeth E. Hwang, Denes Hnisz, Caleb A. Lareau, Linda Ross, Shan Lin, Filemon S. Dela Cruz, Melissa Richardson, Abraham S. Weintraub, Sarah Wang, Amanda Balboni Iniguez, Neekesh V. Dharia, Amy Saur Conway, Amanda L. Robichaud, Benjamin Tanenbaum, John M. Krill-Burger, Francisca Vazquez, Monica Schenone, Jason N. Berman, Andrew L. Kung, Steven A. Carr, Martin J. Aryee, Richard A. Young, Brian D. Crompton, and Kimberly Stegmaier. 2021. ‘STAG2 loss rewires oncogenic and developmental programs to promote metastasis in Ewing sarcoma’, Cancer Cell, 39: 827–44.e10.

Alerasool, N., D. Segal, H. Lee, and M. Taipale. 2020. ‘An efficient KRAB domain for CRISPRi applications in human cells’, Nat Methods, 17: 1093–96.

Alonso-Gil, D., A. Cuadrado, D. Giménez-Llorente, M. Rodríguez-Corsino, and A. Losada. 2023. ‘Different NIPBL requirements of cohesin-STAG1 and cohesin-STAG2’, Nat Commun, 14: 1326.

Arruda, Nicole L., Zachary M. Carico, Megan Justice, Ying Frances Liu, Junjie Zhou, Holden C. Stefan, and Jill M. Dowen. 2020. ‘Distinct and overlapping roles of STAG1 and STAG2 in cohesin localization and gene expression in embryonic stem cells’, Epigenetics & Chromatin, 13: 32.

Ashkin, E. L., Y. J. Tang, H. Xu, K. L. Hung, J. A. Belk, H. Cai, S. S. Lopez, D. N. Dolcen, J. D. Hebert, R. Li, P. A. Ruiz, T. Keal, L. Andrejka, H. Y. Chang, D. A. Petrov, J. R. Dixon, Z. Xu, and M. M. Winslow. 2025. ‘A STAG2-PAXIP1/PAGR1 axis suppresses lung tumorigenesis’, J Exp Med, 222.

Bailey, M. L., D. Tieu, A. Habsid, A. H. Y. Tong, K. Chan, J. Moffat, and P. Hieter. 2021. ‘Paralogous synthetic lethality underlies genetic dependencies of the cancer-mutated gene STAG2’, Life Sci Alliance, 4.

Bailey, Melanie L., Nigel J. O’Neil, Derek M. van Pel, David A. Solomon, Todd Waldman, and Philip Hieter. 2014. ‘Glioblastoma Cells Containing Mutations in the Cohesin Component STAG2 Are Sensitive to PARP Inhibition’, Molecular Cancer Therapeutics, 13: 724–32.

Balbás-Martínez, Cristina, Ana Sagrera, Enrique Carrillo-de-Santa-Pau, Julie Earl, Mirari Márquez, Miguel Vazquez, Eleonora Lapi, Francesc Castro-Giner, Sergi Beltran, Mònica Bayés, Alfredo Carrato, Juan C. Cigudosa, Orlando Domínguez, Marta Gut, Jesús Herranz, Núria Juanpere, Manolis Kogevinas, Xavier Langa, Elena López-Knowles, José A. Lorente, Josep Lloreta, David G. Pisano, Laia Richart, Daniel Rico, Rocío N. Salgado, Adonina Tardón, Stephen Chanock, Simon Heath, Alfonso Valencia, Ana Losada, Ivo Gut, Núria Malats, and Francisco X. Real. 2013. ‘Recurrent inactivation of STAG2 in bladder cancer is not associated with aneuploidy’, Nature Genetics, 45: 1464–69.

Banaszynski, L. A., L. C. Chen, L. A. Maynard-Smith, A. G. Ooi, and T. J. Wandless. 2006. ‘A rapid, reversible, and tunable method to regulate protein function in living cells using synthetic small molecules’, Cell, 126: 995–1004.

Benedict, B., J. J. M. van Schie, A. B. Oostra, J. A. Balk, R. M. F. Wolthuis, H. T. Riele, and J. de Lange. 2020. ‘WAPL-Dependent Repair of Damaged DNA Replication Forks Underlies Oncogene-Induced Loss of Sister Chromatid Cohesion’, Dev Cell, 52: 683–98.e7.

Brough, R., I. Bajrami, R. Vatcheva, R. Natrajan, J. S. Reis-Filho, C. J. Lord, and A. Ashworth. 2012. ‘APRIN is a cell cycle specific BRCA2-interacting protein required for genome integrity and a predictor of outcome after chemotherapy in breast cancer’, Embo j, 31: 1160–76.

Bryant, Helen E., Niklas Schultz, Huw D. Thomas, Kayan M. Parker, Dan Flower, Elena Lopez, Suzanne Kyle, Mark Meuth, Nicola J. Curtin, and Thomas Helleday. 2005. ‘Specific killing of BRCA2-deficient tumours with inhibitors of poly(ADP-ribose) polymerase’, Nature, 434: 913–17.

Casa, V., M. Moronta Gines, E. Gade Gusmao, J. A. Slotman, A. Zirkel, N. Josipovic, E. Oole, IJcken W. F. J. van, A. B. Houtsmuller, A. Papantonis, and K. S. Wendt. 2020. ‘Redundant and specific roles of cohesin STAG subunits in chromatin looping and transcriptional control’, Genome Res, 30: 515–27.

Chung, H. K., C. L. Jacobs, Y. Huo, J. Yang, S. A. Krumm, R. K. Plemper, R. Y. Tsien, and M. Z. Lin. 2015. ‘Tunable and reversible drug control of protein production via a self-excising degron’, Nat Chem Biol, 11: 713–20.

Ciosk, R., M. Shirayama, A. Shevchenko, T. Tanaka, A. Toth, A. Shevchenko, and K. Nasmyth. 2000. ‘Cohesin’s binding to chromosomes depends on a separate complex consisting of Scc2 and Scc4 proteins’, Mol Cell, 5: 243–54.

Cong, K., M. Peng, A. N. Kousholt, W. T. C. Lee, S. Lee, S. Nayak, J. Krais, P. S. VanderVere-Carozza, K. S. Pawelczak, J. Calvo, N. J. Panzarino, J. J. Turchi, N. Johnson, J. Jonkers, E. Rothenberg, and S. B. Cantor. 2021. ‘Replication gaps are a key determinant of PARP inhibitor synthetic lethality with BRCA deficiency’, Mol Cell, 81: 3227.

Couturier, A. M., H. Fleury, A. M. Patenaude, V. L. Bentley, A. Rodrigue, Y. Coulombe, J. Niraj, J. Pauty, J. N. Berman, G. Dellaire, J. M. Di Noia, A. M. Mes-Masson, and J. Y. Masson. 2016. ‘Roles for APRIN (PDS5B) in homologous recombination and in ovarian cancer prediction’, Nucleic Acids Res, 44: 10879–97.

Davidson, I. F., B. Bauer, D. Goetz, W. Tang, G. Wutz, and J. M. Peters. 2019. ‘DNA loop extrusion by human cohesin’, Science, 366: 1338–45.

Delamarre, A., A. Barthe, C. de la Roche Saint-André, P. Luciano, R. Forey, I. Padioleau, M. Skrzypczak, K. Ginalski, V. Géli, P. Pasero, and A. Lengronne. 2020. ‘MRX Increases Chromatin Accessibility at Stalled Replication Forks to Promote Nascent DNA Resection and Cohesin Loading’, Mol Cell, 77: 395–410.e3.

Dobin, Alexander, Carrie A. Davis, Felix Schlesinger, Jorg Drenkow, Chris Zaleski, Sonali Jha, Philippe Batut, Mark Chaisson, and Thomas R. Gingeras. 2013. ‘STAR: ultrafast universal RNA-seq aligner’, Bioinformatics, 29: 15–21.

Farmer, Hannah, Nuala McCabe, Christopher J. Lord, Andrew N. J. Tutt, Damian A. Johnson, Tobias B. Richardson, Manuela Santarosa, Krystyna J. Dillon, Ian Hickson, Charlotte Knights, Niall M. B. Martin, Stephen P. Jackson, Graeme C. M. Smith, and Alan Ashworth. 2005. ‘Targeting the DNA repair defect in BRCA mutant cells as a therapeutic strategy’, Nature, 434: 917–21.

Fellmann, Christof, Thomas Hoffmann, Vaishali Sridhar, Barbara Hopfgartner, Matthias Muhar, Mareike Roth, Dan Yu Lai, Inês A M. Barbosa, Jung Shick Kwon, Yuanzhe Guan, Nishi Sinha, and Johannes Zuber. 2013. ‘An Optimized microRNA Backbone for Effective Single-Copy RNAi’, Cell Reports, 5: 1704–13.

Fingerman, D. F., D. R. O’Leary, A. R. Hansen, T. Tran, B. R. Harris, R. A. DeWeerd, K. E. Hayer, J. Fan, E. Chen, M. Tennakoon, A. Meroni, J. H. Szeto, J. Devenport, D. LaVigne, M. D. Weitzman, O. Shalem, J. Bednarski, A. Vindigni, X. Zhao, and A. M. Green. 2024. ‘The SMC5/6 complex prevents genotoxicity upon APOBEC3A-mediated replication stress’, Embo j, 43: 3240–55.

Fischer, Alexander, Benjamín Hernández-Rodríguez, Roger Mulet-Lazaro, Margit Nuetzel, Fabian Hölzl, Stanley van Herk, François G. Kavelaars, Hanna Stanewsky, Ute Ackermann, Amadou H. Niang, Noelia Diaz, Edith Reuschel, Nicholas Strieder, Inmaculada Hernández-López, Peter J. M. Valk, Juan M. Vaquerizas, Michael Rehli, Ruud Delwel, and Claudia Gebhard. 2024. ‘STAG2 mutations reshape the cohesin-structured spatial chromatin architecture to drive gene regulation in acute myeloid leukemia’, Cell Reports, 43: 114498.

Frattini, C., S. Villa-Hernández, G. Pellicanò, R. Jossen, Y. Katou, K. Shirahige, and R. Bermejo. 2017. ‘Cohesin Ubiquitylation and Mobilization Facilitate Stalled Replication Fork Dynamics’, Mol Cell, 68: 758–72.e4.

Frederick, M. I., E. Fyle, A. Clouvel, D. Abdesselam, and S. Hassan. 2025. ‘Targeting FEN1/EXO1 to enhance efficacy of PARP inhibition in triple-negative breast cancer’, Transl Oncol, 54: 102337.

Galeev, R., A. Baudet, P. Kumar, A. Rundberg Nilsson, B. Nilsson, S. Soneji, T. Törngren, Å Borg, A. Kvist, and J. Larsson. 2016. ‘Genome-wide RNAi Screen Identifies Cohesin Genes as Modifiers of Renewal and Differentiation in Human HSCs’, Cell Rep, 14: 2988–3000.

Gopal, A. A., B. Fernandez, J. Delano, R. Weissleder, and J. M. Dubach. 2024. ‘PARP trapping is governed by the PARP inhibitor dissociation rate constant’, Cell Chem Biol, 31: 1373–82.e10.

Hage Chehade, C., G. Gebrael, N. Sayegh, Z. I. Ozay, A. Narang, T. Crispino, T. Golan, J. K. Litton, U. Swami, K. N. Moore, and N. Agarwal. 2025. ‘A pan-tumor review of the role of poly(adenosine diphosphate ribose) polymerase inhibitors’, CA Cancer J Clin, 75: 141–67.

Herzog, M., F. Puddu, J. Coates, N. Geisler, J. V. Forment, and S. P. Jackson. 2018. ‘Detection of functional protein domains by unbiased genome-wide forward genetic screening’, Sci Rep, 8: 6161.

Hill, V. K., J. S. Kim, and T. Waldman. 2016. ‘Cohesin mutations in human cancer’, Biochim Biophys Acta, 1866: 1–11.

Hou, W., Y. Li, J. Zhang, Y. Xia, X. Wang, H. Chen, and H. Lou. 2022. ‘Cohesin in DNA damage response and double-strand break repair’, Crit Rev Biochem Mol Biol, 57: 333–50.

Hu, B., T. Itoh, A. Mishra, Y. Katoh, K. L. Chan, W. Upcher, C. Godlee, M. B. Roig, K. Shirahige, and K. Nasmyth. 2011. ‘ATP hydrolysis is required for relocating cohesin from sites occupied by its Scc2/4 loading complex’, Curr Biol, 21: 12–24.

Jacobs, C. L., R. K. Badiee, and M. Z. Lin. 2018. ‘StaPLs: versatile genetically encoded modules for engineering drug-inducible proteins’, Nat Methods, 15: 523–26.

Jahjah, Tiya, Jenny K. Singh, Vanesa Gottifredi, and Annabel Quinet. 2024. ‘Tolerating DNA damage by repriming: Gap filling in the spotlight’, DNA Repair, 142: 103758.

Kawale, A. S., X. Ran, P. S. Patel, S. Saxena, M. S. Lawrence, and L. Zou. 2024. ‘APOBEC3A induces DNA gaps through PRIMPOL and confers gap-associated therapeutic vulnerability’, Sci Adv, 10: eadk2771.

Kim, Jung-Sik, Xiaoyuan He, Bernardo Orr, Gordana Wutz, Victoria Hill, Jan-Michael Peters, Duane A. Compton, and Todd Waldman. 2016. ‘Intact Cohesion, Anaphase, and Chromosome Segregation in Human Cells Harboring Tumor-Derived Mutations in STAG2’, PLOS Genetics, 12: e1005865.

Kojic, A., A. Cuadrado, M. De Koninck, D. Giménez-Llorente, M. Rodríguez-Corsino, G. Gómez-López, F. Le Dily, M. A. Marti-Renom, and A. Losada. 2018. ‘Distinct roles of cohesin-SA1 and cohesin-SA2 in 3D chromosome organization’, Nat Struct Mol Biol, 25: 496–504.

Konermann, Silvana, Peter Lotfy, Nicholas J. Brideau, Jennifer Oki, Maxim N. Shokhirev, and Patrick D. Hsu. 2018. ‘Transcriptome Engineering with RNA-Targeting Type VI-D CRISPR Effectors’, Cell, 173: 665–76.e14.

Kukolj, E., T. Kaufmann, A. E. Dick, R. Zeillinger, D. W. Gerlich, and D. Slade. 2017. ‘PARP inhibition causes premature loss of cohesion in cancer cells’, Oncotarget, 8: 103931–51.

Li, Y., J. H. I. Haarhuis, Á Sedeño Cacciatore, R. Oldenkamp, M. S. van Ruiten, L. Willems, H. Teunissen, K. W. Muir, E. de Wit, B. D. Rowland, and D. Panne. 2020. ‘The structural basis for cohesin-CTCF-anchored loops’, Nature, 578: 472–76.

Li, Y., K. W. Muir, M. W. Bowler, J. Metz, C. H. Haering, and D. Panne. 2018. ‘Structural basis for Scc3-dependent cohesin recruitment to chromatin’, Elife, 7.

Litwin, Ireneusz, Ewa Pilarczyk, and Robert Wysocki. 2018. ‘The Emerging Role of Cohesin in the DNA Damage Response’, Genes, 9: 581.

Losada, A., T. Yokochi, R. Kobayashi, and T. Hirano. 2000. ‘Identification and characterization of SA/Scc3p subunits in the Xenopus and human cohesin complexes’, J Cell Biol, 150: 405–16.

Love, Michael I., Wolfgang Huber, and Simon Anders. 2014. ‘Moderated estimation of fold change and dispersion for RNA-seq data with DESeq2’, Genome Biology, 15: 550.

Marsischky, G. T., B. A. Wilson, and R. J. Collier. 1995. ‘Role of glutamic acid 988 of human poly-ADP-ribose polymerase in polymer formation. Evidence for active site similarities to the ADP-ribosylating toxins’, J Biol Chem, 270: 3247–54.

Mondal, G., M. Stevers, B. Goode, A. Ashworth, and D. A. Solomon. 2019. ‘A requirement for STAG2 in replication fork progression creates a targetable synthetic lethality in cohesin-mutant cancers’, Nat Commun, 10: 1686.

Morales, C., M. Ruiz-Torres, S. Rodríguez-Acebes, V. Lafarga, M. Rodríguez-Corsino, D. Megías, D. A. Cisneros, J. M. Peters, J. Méndez, and A. Losada. 2020. ‘PDS5 proteins are required for proper cohesin dynamics and participate in replication fork protection’, J Biol Chem, 295: 146–57.

Moronta Gines, M., M. W. Wessels, V. Casa, T. van Staveren, A. Hof, W. K. Chung, M. Willems, A. Sandestig, I. Huening, P. Turnpenny, M. Lefebvre, I. Parenti, F. J. Kaiser, J. Demmers, W. F. J. van Ijcken, and K. S. Wendt. 2025. ‘STAG2-truncating variants reveal a mosaic STAG2 inactivation pattern and compensatory mechanisms involving cohesin complex remodeling’, iScience, 28: 114195.

Mullenders, J., B. Aranda-Orgilles, P. Lhoumaud, M. Keller, J. Pae, K. Wang, C. Kayembe, P. P. Rocha, R. Raviram, Y. Gong, P. K. Premsrirut, A. Tsirigos, R. Bonneau, J. A. Skok, L. Cimmino, D. Hoehn, and I. Aifantis. 2015. ‘Cohesin loss alters adult hematopoietic stem cell homeostasis, leading to myeloproliferative neoplasms’, J Exp Med, 212: 1833–50.

Murai, Junko, Shar-yin N. Huang, Benu Brata Das, Amelie Renaud, Yiping Zhang, James H. Doroshow, Jiuping Ji, Shunichi Takeda, and Yves Pommier. 2012. ‘Trapping of PARP1 and PARP2 by Clinical PARP Inhibitors’, Cancer Research, 72: 5588–99.

Murai, Junko, Shar-Yin N. Huang, Amèlie Renaud, Yiping Zhang, Jiuping Ji, Shunichi Takeda, Joel Morris, Beverly Teicher, James H. Doroshow, and Yves Pommier. 2014. ‘Stereospecific PARP Trapping by BMN 673 and Comparison with Olaparib and Rucaparib’, Molecular Cancer Therapeutics, 13: 433–43.

Murayama, Y., and F. Uhlmann. 2014. ‘Biochemical reconstitution of topological DNA binding by the cohesin ring’, Nature, 505: 367–71.

Nishiyama, T. 2019. ‘Cohesion and cohesin-dependent chromatin organization’, Curr Opin Cell Biol, 58: 8–14.

Pai Bellare, G., K. Kundu, P. Dey, K. T. Philip, N. Chauhan, M. Sharma, S. K. Rajput, and B. S. Patro. 2025. ‘Targeting Replication Fork Processing Synergizes with PARP Inhibition to Potentiate Lethality in Homologous Recombination Proficient Ovarian Cancers’, Adv Sci (Weinh), 12: e2410718.

Perea-Resa, C., L. Wattendorf, S. Marzouk, and M. D. Blower. 2021. ‘Cohesin: behind dynamic genome topology and gene expression reprogramming’, Trends Cell Biol, 31: 760–73.

Peters, J. M., and T. Nishiyama. 2012. ‘Sister chromatid cohesion’, Cold Spring Harb Perspect Biol, 4.

Pettitt, S. J., F. L. Rehman, I. Bajrami, R. Brough, F. Wallberg, I. Kozarewa, K. Fenwick, I. Assiotis, L. Chen, J. Campbell, C. J. Lord, and A. Ashworth. 2013. ‘A genetic screen using the PiggyBac transposon in haploid cells identifies Parp1 as a mediator of olaparib toxicity’, PLoS One, 8: e61520.

Pommier, Y., M. J. O’Connor, and J. de Bono. 2016. ‘Laying a trap to kill cancer cells: PARP inhibitors and their mechanisms of action’, Sci Transl Med, 8: 362ps17.

Replogle, Joseph M., Jessica L. Bonnar, Angela N. Pogson, Christina R. Liem, Nolan K. Maier, Yufang Ding, Baylee J. Russell, Xingren Wang, Kun Leng, Alina Guna, Thomas M. Norman, Ryan A. Pak, Daniel M. Ramos, Michael E. Ward, Luke A. Gilbert, Martin Kampmann, Jonathan S. Weissman, and Marco Jost. 2022. ‘Maximizing CRISPRi efficacy and accessibility with dual-sgRNA libraries and optimal effectors’, Elife, 11: e81856.

Richart, Laia, Eleonora Lapi, Vera Pancaldi, Mirabai Cuenca-Ardura, Enrique Carrillo-de-Santa Pau, Miguel Madrid-Mencía, Hélène Neyret-Kahn, François Radvanyi, Juan Antonio Rodríguez, Yasmina Cuartero, François Serra, François Le Dily, Alfonso Valencia, Marc A Marti-Renom, and Francisco X Real. 2021. ‘STAG2 loss-of-function affects short-range genomic contacts and modulates the basal-luminal transcriptional program of bladder cancer cells’, Nucleic Acids Research, 49: 11005–21.

Rittenhouse, N. L., and J. M. Dowen. 2024. ‘Cohesin regulation and roles in chromosome structure and function’, Curr Opin Genet Dev, 85: 102159.

Rolli, V., M. O’Farrell, J. Ménissier-de Murcia, and G. de Murcia. 1997. ‘Random mutagenesis of the poly(ADP-ribose) polymerase catalytic domain reveals amino acids involved in polymer branching’, Biochemistry, 36: 12147–54.

Romero-Pérez, L., D. Surdez, E. Brunet, O. Delattre, and T. G. P. Grünewald. 2019. ‘STAG Mutations in Cancer’, Trends Cancer, 5: 506–20.

Rowley, M. J., and V. G. Corces. 2018. ‘Organizational principles of 3D genome architecture’, Nat Rev Genet, 19: 789–800.

Saha, L. K., Y. Murai, S. Saha, U. Jo, M. Tsuda, S. Takeda, and Y. Pommier. 2021. ‘Replication-dependent cytotoxicity and Spartan-mediated repair of trapped PARP1-DNA complexes’, Nucleic Acids Res, 49: 10493–506.

Sanjana, N. E., O. Shalem, and F. Zhang. 2014. ‘Improved vectors and genome-wide libraries for CRISPR screening’, Nat Methods, 11: 783–84.

Sasca, D., H. Yun, G. Giotopoulos, J. Szybinski, T. Evan, N. K. Wilson, M. Gerstung, P. Gallipoli, A. R. Green, R. Hills, N. Russell, C. S. Osborne, E. Papaemmanuil, B. Göttgens, P. Campbell, and B. J. P. Huntly. 2019. ‘Cohesin-dependent regulation of gene expression during differentiation is lost in cohesin-mutated myeloid malignancies’, Blood, 134: 2195–208.

Scott, Julia S., Loubna Al Ayadi, Emmanouela Epeslidou, Roan H. van Scheppingen, Anna Mukha, Lucas J. T. Kaaij, Catrin Lutz, and Stefan Prekovic. 2025. ‘Emerging roles of cohesin-STAG2 in cancer’, Oncogene, 44: 277–87.

Shao, Z., B. J. Lee, É Rouleau-Turcotte, M. F. Langelier, X. Lin, V. M. Estes, J. M. Pascal, and S. Zha. 2020. ‘Clinical PARP inhibitors do not abrogate PARP1 exchange at DNA damage sites in vivo’, Nucleic Acids Res, 48: 9694–709.

Shi, Peiguo, Michael R. Murphy, Alexis O. Aparicio, Jordan S. Kesner, Zhou Fang, Ziheng Chen, Aditi Trehan, Yang Guo, and Xuebing Wu. 2023. ‘Collateral activity of the CRISPR/RfxCas13d system in human cells’, Communications Biology, 6: 334.

Shi, Zhubing, Haishan Gao, Xiao-chen Bai, and Hongtao Yu. 2020. ‘Cryo-EM structure of the human cohesin-NIPBL-DNA complex’, Science, 368: 1454–59.

Simoneau, Antoine, Rosalinda Xiong, and Lee Zou. 2021. ‘The trans cell cycle effects of PARP inhibitors underlie their selectivity toward BRCA1/2-deficient cells’, Genes & Development, 35: 1271–89.

Smith, J. S., K. M. Lappin, S. G. Craig, F. G. Liberante, C. M. Crean, S. S. McDade, A. Thompson, K. I. Mills, and K. I. Savage. 2020. ‘Chronic loss of STAG2 leads to altered chromatin structure contributing to de-regulated transcription in AML’, J Transl Med, 18: 339.

Solomon, D. A., T. Kim, L. A. Diaz-Martinez, J. Fair, A. G. Elkahloun, B. T. Harris, J. A. Toretsky, S. A. Rosenberg, N. Shukla, M. Ladanyi, Y. Samuels, C. D. James, H. Yu, J. S. Kim, and T. Waldman. 2011. ‘Mutational inactivation of STAG2 causes aneuploidy in human cancer’, Science, 333: 1039–43.

Srinivasan, Madhusudhan, Marco Fumasoni, Naomi J. Petela, Andrew Murray, and Kim A. Nasmyth. 2020. ‘Cohesion is established during DNA replication utilising chromosome associated cohesin rings as well as those loaded de novo onto nascent DNAs’, Elife, 9: e56611.

Sumara, I., E. Vorlaufer, C. Gieffers, B. H. Peters, and J. M. Peters. 2000. ‘Characterization of vertebrate cohesin complexes and their regulation in prophase’, J Cell Biol, 151: 749–62.

Surdez, Didier, Sakina Zaidi, Sandrine Grossetête, Karine Laud-Duval, Anna Sole Ferre, Lieke Mous, Thomas Vourc’h, Franck Tirode, Gaelle Pierron, Virginie Raynal, Sylvain Baulande, Erika Brunet, Véronique Hill, and Olivier Delattre. 2021. ‘STAG2 mutations alter CTCF-anchored loop extrusion, reduce cis-regulatory interactions and EWSR1-FLI1 activity in Ewing sarcoma’, Cancer Cell, 39: 810–26.e9.

Teloni, Federico, Zsuzsanna Takacs, Michael Mitter, Christoph C. H. Langer, Inès Prlesi, Thomas L. Steinacker, Vincent P. Reuter, Dmitry Mylarshchikov, and Daniel W. Gerlich. 2025. ‘Cohesin guides homology search during DNA repair using loops and sister chromatid linkages’, Science, 390: eadw0566.

Tirman, S., A. Quinet, M. Wood, A. Meroni, E. Cybulla, J. Jackson, S. Pegoraro, A. Simoneau, L. Zou, and A. Vindigni. 2021. ‘Temporally distinct post-replicative repair mechanisms fill PRIMPOL-dependent ssDNA gaps in human cells’, Mol Cell, 81: 4026–40.e8.

Tittel-Elmer, M., A. Lengronne, M. B. Davidson, J. Bacal, P. François, M. Hohl, J. H. J. Petrini, P. Pasero, and J. A. Cobb. 2012. ‘Cohesin association to replication sites depends on rad50 and promotes fork restart’, Mol Cell, 48: 98–108.

Tothova, Z., A. L. Valton, R. A. Gorelov, M. Vallurupalli, J. M. Krill-Burger, A. Holmes, C. C. Landers, J. E. Haydu, E. Malolepsza, C. Hartigan, M. Donahue, K. D. Popova, S. Koochaki, S. V. Venev, J. Rivera, E. Chen, K. Lage, M. Schenone, A. D. D’Andrea, S. A. Carr, E. A. Morgan, J. Dekker, and B. L. Ebert. 2021. ‘Cohesin mutations alter DNA damage repair and chromatin structure and create therapeutic vulnerabilities in MDS/AML’, JCI Insight, 6.

Trotter, Eleanor Wendy, and Iain Michael Hagan. 2020. ‘Release from cell cycle arrest with Cdk4/6 inhibitors generates highly synchronized cell cycle progression in human cell culture’, Open Biology, 10: 200200.

Vaitsiankova, A., K. Burdova, M. Sobol, A. Gautam, O. Benada, H. Hanzlikova, and K. W. Caldecott. 2022. ‘PARP inhibition impedes the maturation of nascent DNA strands during DNA replication’, Nat Struct Mol Biol, 29: 329–38.

van der Lelij, P., S. Lieb, J. Jude, G. Wutz, C. P. Santos, K. Falkenberg, A. Schlattl, J. Ban, R. Schwentner, T. Hoffmann, H. Kovar, F. X. Real, T. Waldman, M. A. Pearson, N. Kraut, J. M. Peters, J. Zuber, and M. Petronczki. 2017. ‘Synthetic lethality between the cohesin subunits STAG1 and STAG2 in diverse cancer contexts’, Elife, 6.

van der Weegen, Y., K. de Lint, D. van den Heuvel, Y. Nakazawa, T. E. T. Mevissen, J. J. M. van Schie, M. San Martin Alonso, D. E. C. Boer, R. González-Prieto, I. V. Narayanan, N. H. M. Klaassen, A. P. Wondergem, K. Roohollahi, J. C. Dorsman, Y. Hara, A. C. O. Vertegaal, J. de Lange, J. C. Walter, S. M. Noordermeer, M. Ljungman, T. Ogi, R. M. F. Wolthuis, and M. S. Luijsterburg. 2021. ‘ELOF1 is a transcription-coupled DNA repair factor that directs RNA polymerase II ubiquitylation’, Nat Cell Biol, 23: 595–607.

van Schie, J. J. M., K. de Lint, T. M. Molenaar, M. Moronta Gines, J. A. Balk, M. A. Rooimans, K. Roohollahi, G. M. Pai, L. Borghuis, A. R. Ramadhin, F. Corazza, J. C. Dorsman, K. S. Wendt, R. M. F. Wolthuis, and J. de Lange. 2023. ‘CRISPR screens in sister chromatid cohesion defective cells reveal PAXIP1-PAGR1 as regulator of chromatin association of cohesin’, Nucleic Acids Res, 51: 9594–609.

van Schie, J. J. M., A. Faramarz, J. A. Balk, G. S. Stewart, E. Cantelli, A. B. Oostra, M. A. Rooimans, J. L. Parish, C. de Almeida Estéves, K. Dumic, I. Barisic, K. E. M. Diderich, M. A. van Slegtenhorst, M. Mahtab, F. M. Pisani, H. Te Riele, N. Ameziane, R. M. F. Wolthuis, and J. de Lange. 2020. ‘Warsaw Breakage Syndrome associated DDX11 helicase resolves G-quadruplex structures to support sister chromatid cohesion’, Nat Commun, 11: 4287.

van Schie, Janne JM, Klaas de Lint, Govind M Pai, Martin A Rooimans, Rob MF Wolthuis, and Job de Lange. 2023. ‘MMS22L-TONSL functions in sister chromatid cohesion in a pathway parallel to DSCC1-RFC’, Life Science Alliance, 6: e202201596.

Viny, A. D., R. L. Bowman, Y. Liu, V. P. Lavallée, S. E. Eisman, W. Xiao, B. H. Durham, A. Navitski, J. Park, S. Braunstein, B. Alija, A. Karzai, I. S. Csete, M. Witkin, E. Azizi, T. Baslan, C. J. Ott, D. Pe’er, J. Dekker, R. Koche, and R. L. Levine. 2019. ‘Cohesin Members Stag1 and Stag2 Display Distinct Roles in Chromatin Accessibility and Topological Control of HSC Self-Renewal and Differentiation’, Cell Stem Cell, 25: 682–96.e8.

Waldman, Todd. 2020. ‘Emerging themes in cohesin cancer biology’, Nature Reviews Cancer, 20: 504– 15.

Wutz, G., R. Ladurner, B. G. St Hilaire, R. R. Stocsits, K. Nagasaka, B. Pignard, A. Sanborn, W. Tang, C. Várnai, M. P. Ivanov, S. Schoenfelder, P. van der Lelij, X. Huang, G. Dürnberger, E. Roitinger, K. Mechtler, I. F. Davidson, P. Fraser, E. Lieberman-Aiden, and J. M. Peters. 2020. ‘ESCO1 and CTCF enable formation of long chromatin loops by protecting cohesin(STAG1) from WAPL’, Elife, 9.

Yatskevich, S., J. Rhodes, and K. Nasmyth. 2019. ‘Organization of Chromosomal DNA by SMC Complexes’, Annu Rev Genet, 53: 445–82.

Zeng, Y., O. Arisa, C. J. Peer, A. Fojo, and W. D. Figg. 2024. ‘PARP inhibitors: A review of the pharmacology, pharmacokinetics, and pharmacogenetics’, Semin Oncol, 51: 19–24.

Zhou, J., R. C. Nie, Z. P. He, X. X. Cai, J. W. Chen, W. P. Lin, Y. X. Yin, Z. C. Xiang, T. C. Zhu, J. J. Xie, Y. C. Zhang, X. Wang, P. Lin, D. Xie, A. D. D’Andrea, and M. Y. Cai. 2023. ‘STAG2 Regulates Homologous Recombination Repair and Sensitivity to ATM Inhibition’, Adv Sci (Weinh), 10: e2302494.

