## Supplementary Figures 1-15 for "Residual cohesin function and PARP1 levels determine PARP inhibitor sensitivity in STAG2-deficient cancers"

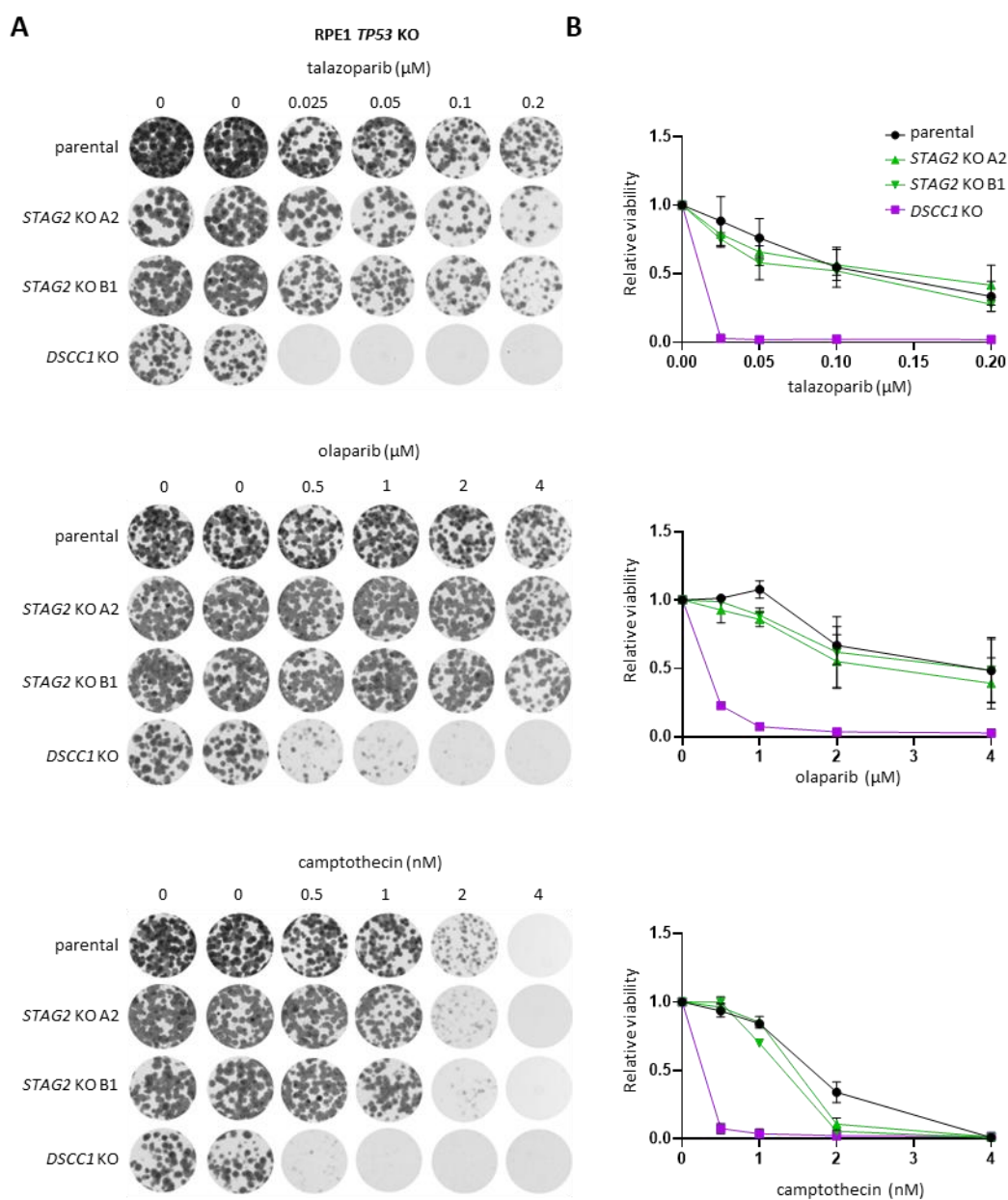

**Supplementary Figure 1. Clonogenic survival assays for STAG2 KO RPE1 p53KO cells. (A)** STAG2 and control DSCC1 KO cells were grown for 14 days with indicated drugs and surviving colonies were stained with crystal violet. **(B)** Quantification of colony area coverage for clonogenic survival assays. Error bars are SD for three independent experiments.

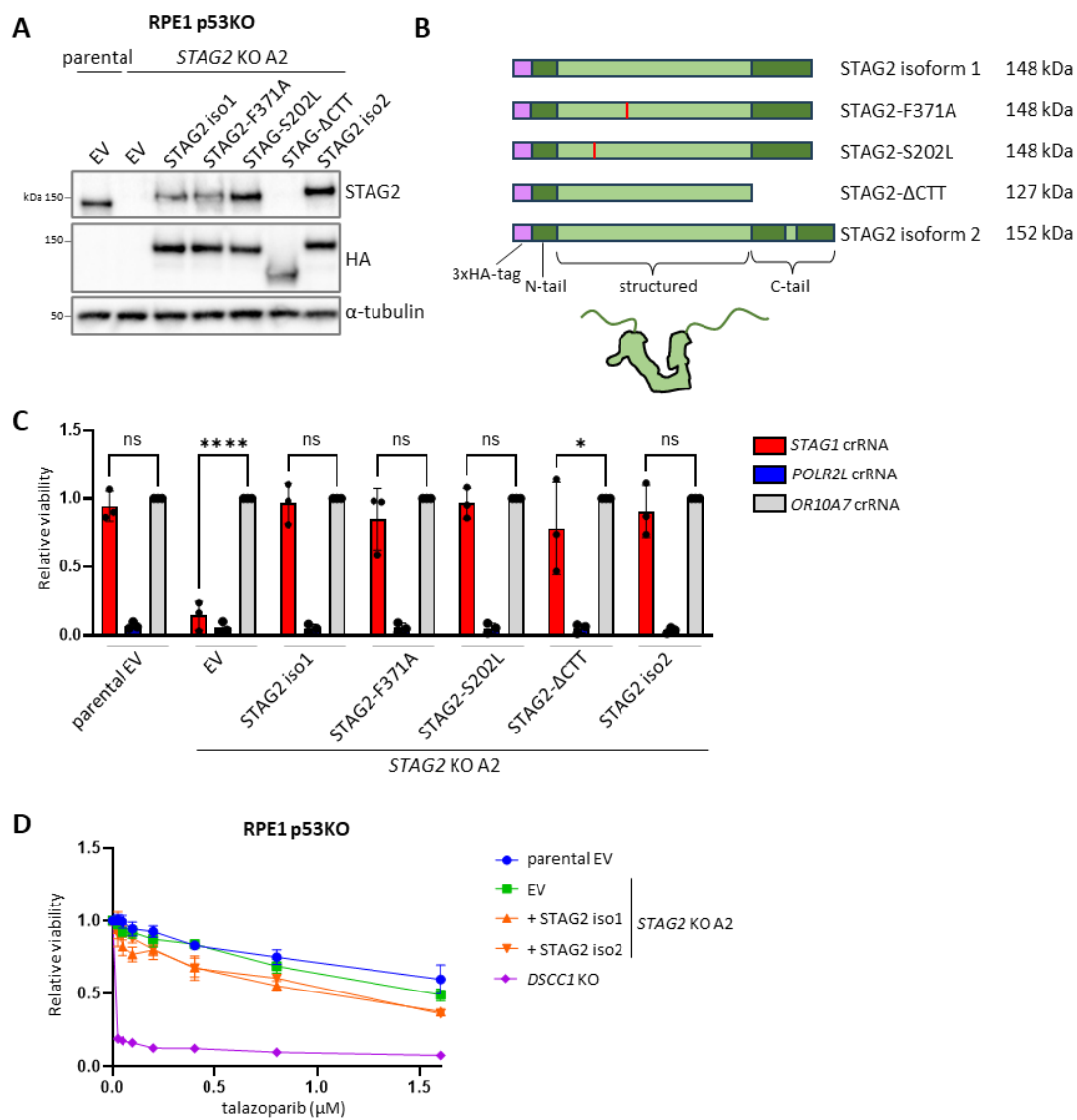

**Supplementary Figure 2. Confirmation of *STAG2* KO status in RPE1 p53KO cells by *STAG1-STAG2* synthetic lethality.** (A) Whole-cell extract western blot of STAG2 ectopic rescue constructs delivered by stable lentiviral transduction. The epitope for the STAG2 antibody is located in the C-terminal region causing the ΔCTT mutant not to appear on the STAG2 blot. (B) Schematic of STAG2 rescue constructs with N-terminal HA tag. The indicated protein weights are including 3xHA-tag. (C) Viability of RPE1 p53KO cells after acute *STAG1* KO. RPE1 p53KO cells expressing dox inducible Cas9 were transfected with tracrRNA plus crRNAs targeting *STAG1*, *POLR2L* (an essential gene), or *OR10A7* (non-essential gene) and cell viability was measured after 9 days. Error bars represent the SD of three independent experiments. Statistical significance was assessed using two-way ANOVA followed by Dunnett's multiple comparisons test. (D) Talazoparib sensitivity of RPE1 p53KO *STAG2* KO cells complemented with STAG2 rescue constructs. Error bars represent SEM for three independent experiments.

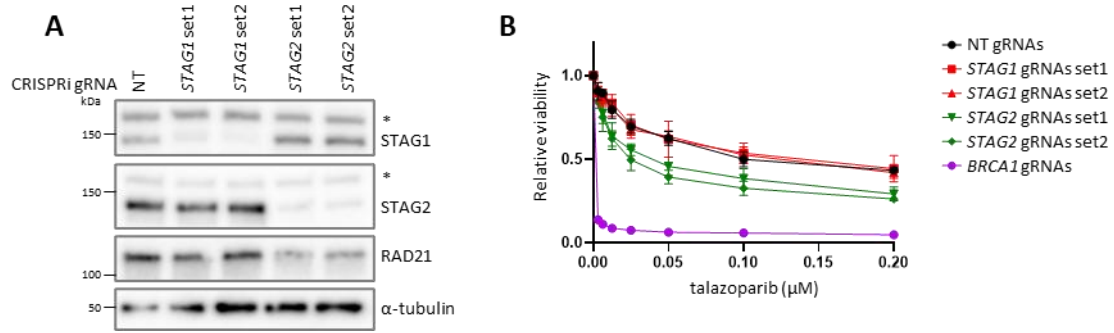

**Supplementary Figure 3. STAG2 KD in K562 using CRISPRi moderately sensitizes cells to talazoparib.**

**(A)** Whole-cell extract western blot after CRISPRi in K562 using simultaneous expression of two gRNAs per gene. Two independent sets of gRNAs were used for each gene. **(B)** Talazoparib sensitivity in K562 CRISPRi cells. Knockdown was induced with dox 48h prior to seeding cells for drug assays. Cells were treated with talazoparib for 6 days. Error bars represent SEM for three independent experiments.

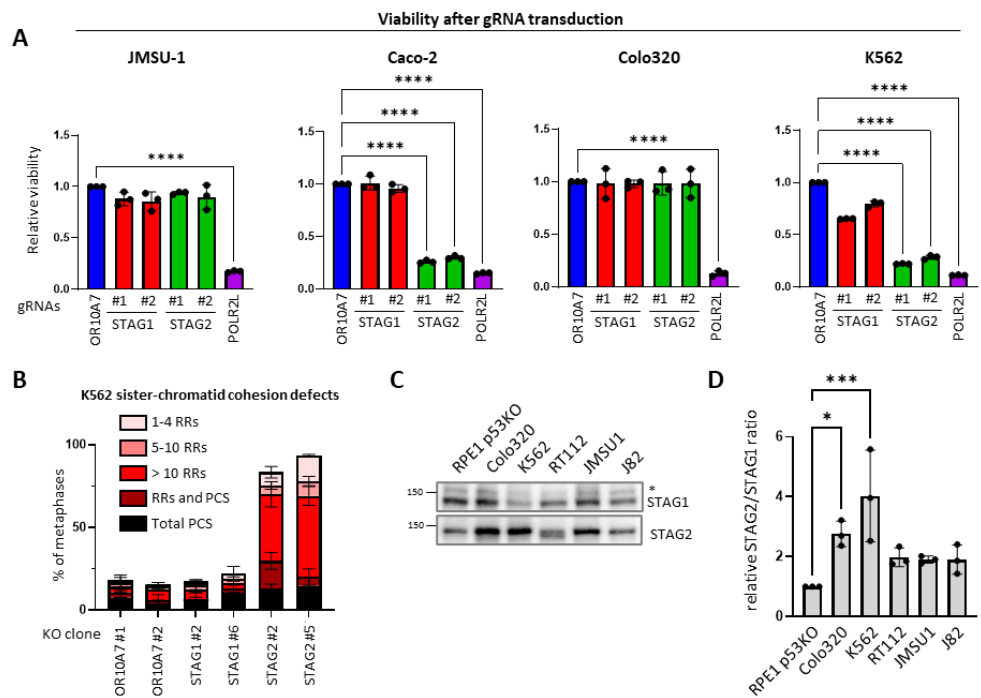

**Supplementary Figure 4. Effect of *STAG2* loss on viability of JMSU-1, Caco-2, Colo320, and K562. (A)** Cas9 expressing cell lines were transduced with vectors encoding two gRNAs against the same gene. Two independent sets of gRNAs were used for *STAG1* and *STAG2*. Transduced cells were selected with puromycin for 3 days and viability was measured 9 days after transduction. **(B)** Sister chromatid cohesion analysis in *STAG1* and *STAG2* KO K562 cells. RR indicates rail-road chromosomes, PCS is premature chromatid separation. Error bars are SD for three independent experiments. **(C)** Western blot showing relative STAG1 and STAG2 levels in different cell lines. **(D)** Quantification of STAG2/STAG1 ratios measured by western blot. For A and D, error bars represent the SD of three independent experiments. Statistical significance was assessed using one-way ANOVA followed by Dunnett's multiple comparisons test.

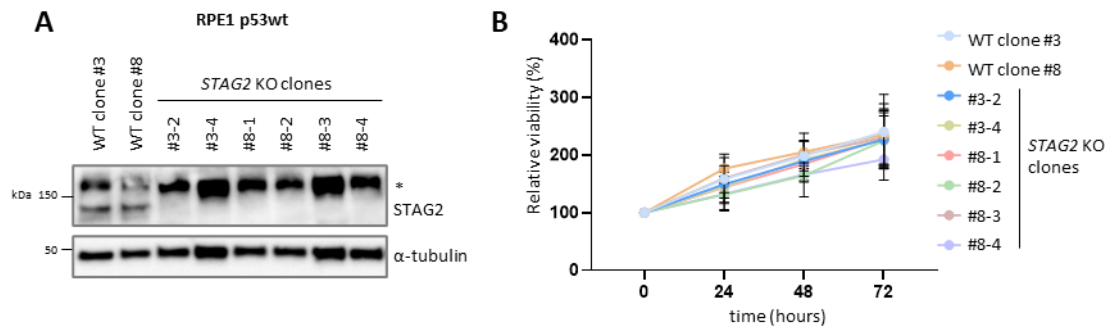

**Supplementary Figure 5. *STAG2* KO has no effect on viability in p53wt RPE1 cells. (A)** Western blot showing STAG2 loss in KO clones derived from two RPE1 p53wt cell lines with dox inducible Cas9. **(B)** Growth assay for *STAG2* KO RPE1 p53wt cell lines. Error bars represent SD for four independent experiments.

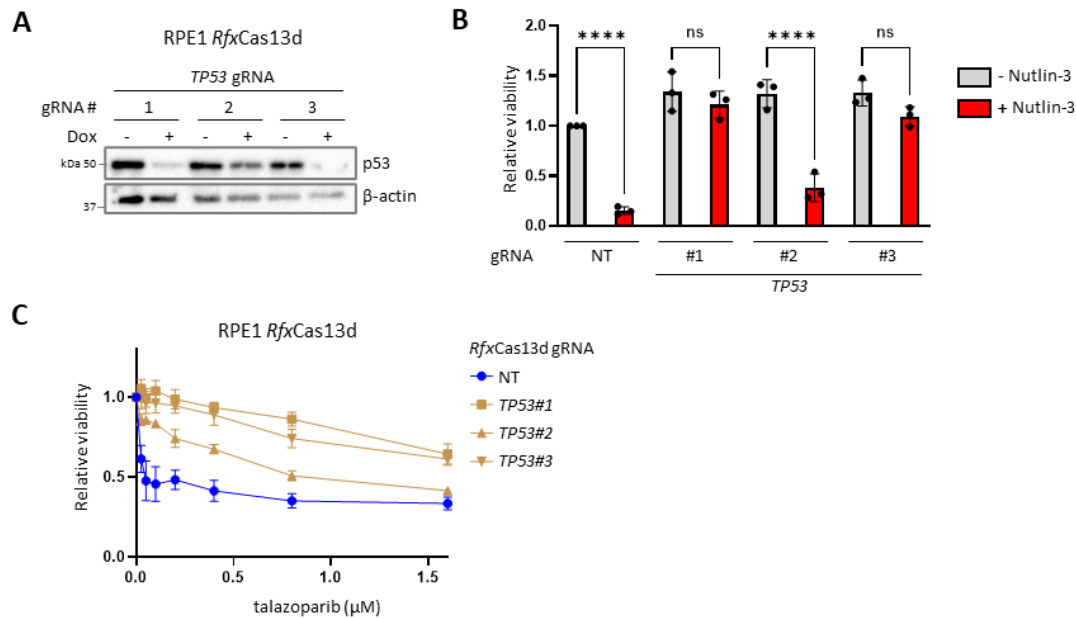

**Supplementary Figure 6. TP53 mRNA depletion makes RPE1 cells resistant to talazoparib. (A)** Western blot showing p53 depletion after *RfxCas13d* mediated mRNA knockdown in RPE1 cells. **(B)** Cell viability was measured after treatment with 5 μM Nutlin-3 for 5 days to show functional loss of the p53 pathway. Error bars represent the SD of three independent experiments. Statistical significance was assessed using one-way ANOVA followed by Dunnett's multiple comparisons test. **(C)** Cell viability assays following treatment with talazoparib for 5 days. Error bars represent the SEM for three independent experiments.

**A**

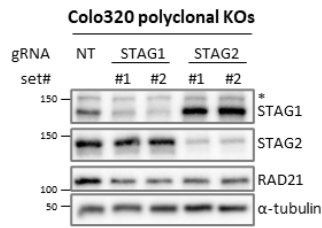

**B**

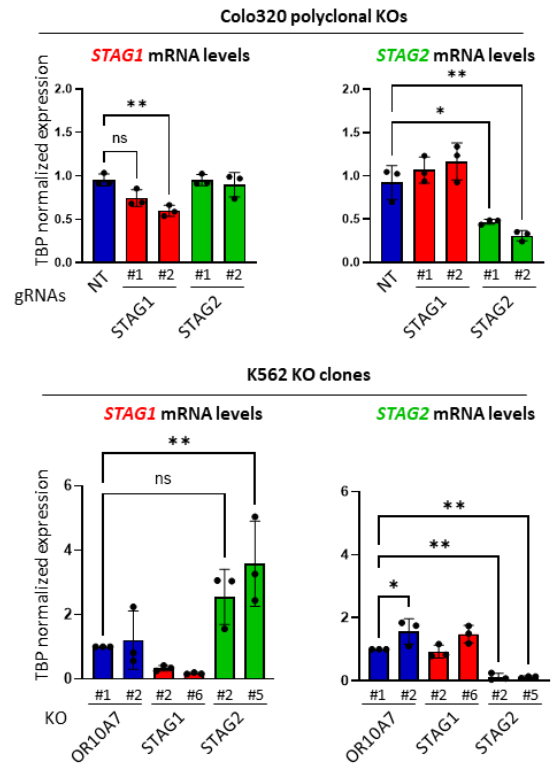

**Supplementary Figure 7. Polyclonal *STAG2* KO Colo320 cells have higher *STAG1* protein levels. (A)** Western blot of Colo320 cells with dox inducible Cas9 transduced with dual gRNAs against *STAG1* or *STAG2*. \* indicates a non-specific band. **(B)** RNA levels of *STAG1* and *STAG2* measured by RT-qPCR in polyclonal *STAG1/2* KO Colo320 cells and monoclonal *STAG1/2* KO K562 cells. Error bars represent the SD of three independent experiments. Statistical significance was assessed using one-way ANOVA followed by Dunnett's multiple comparisons test.

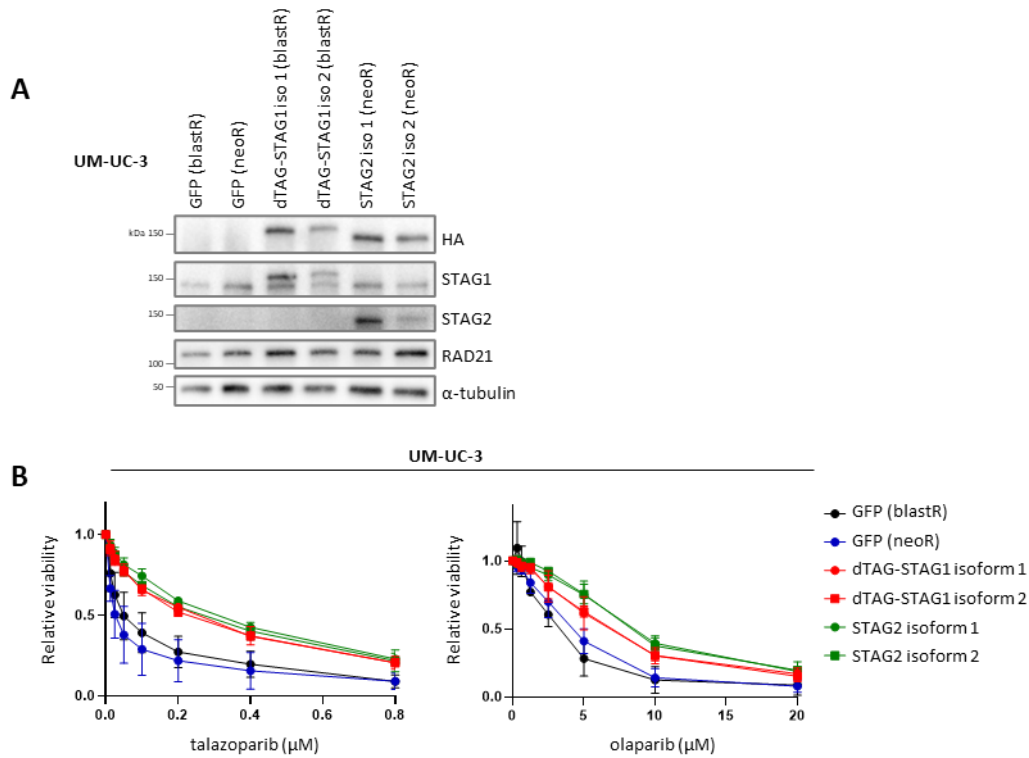

**Supplementary Figure 8. Ectopic expression of either STAG1 or STAG2 desensitizes UM-UC-3 cells to talazoparib. (A)** Western blot of UM-UC-3 cells stably expressing both isoforms of STAG1 or STAG2 with HA tag. Cells transduced with STAG1 were selected using neomycin resistance (neoR), STAG2 with blasticidin resistance (blastR). Note that the epitope recognized by the STAG2 antibody lies within the region altered by alternative splicing that generates STAG2 isoform 2 through inclusion of an additional exon. The antibody therefore likely recognizes both isoforms with different affinity (unlike the HA antibody). **(B)** PARPi sensitivity of STAG1/2 complemented UM-UC-3 cells. Cells were treated with PARPi for 5 days. Error bars represent SEM for three independent experiments.

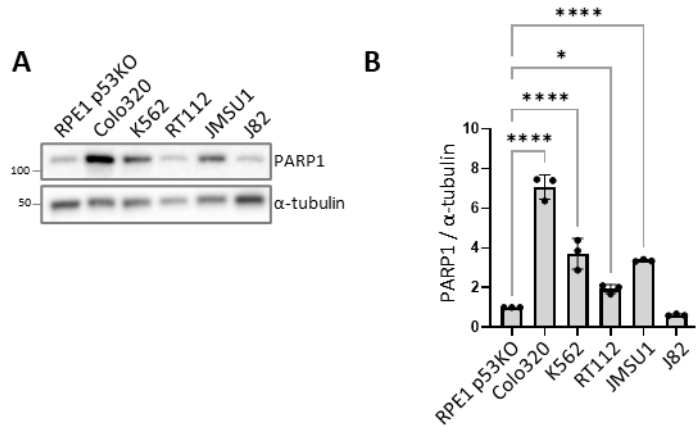

**Supplementary Figure 9. PARP1 protein levels vary among cell lines. (A)** Western blot showing relative PARP1 levels in different cell lines. **(B)** Quantification of PARP1 levels from western blot. Error bars represent the SD of three independent experiments. Statistical significance was assessed using one-way ANOVA followed by Dunnett's multiple comparisons test.

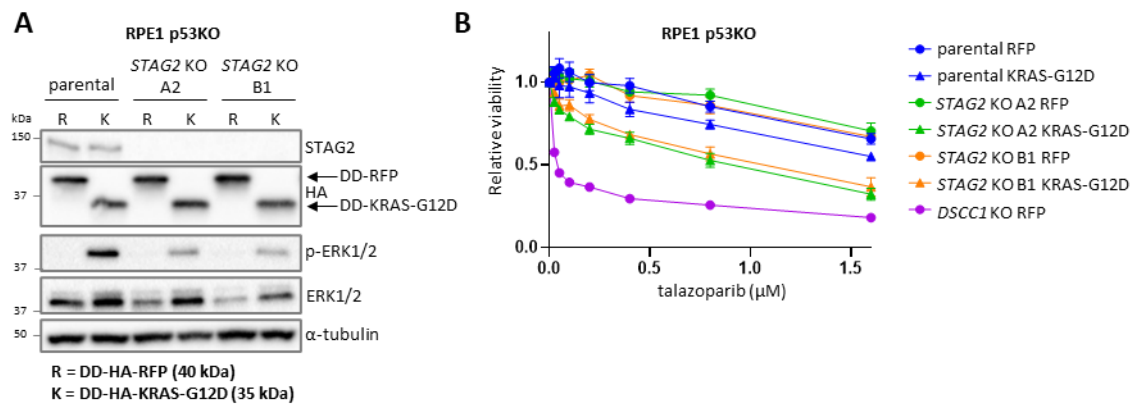

**Supplementary Figure 10. KRAS-G12D overexpression moderately sensitizes RPE1 STAG2 KO cells to talazoparib. (A)** Western blot showing Shield-1 induced overexpression of DD-KRAS-G12D in wt and STAG2 KO RPE1 p53KO cells. Phospho-ERK1/2 (p-ERK1/2) is used as a marker of (hyper)active KRAS signaling. **(B)** Cells were treated with 0.5 μM Shield-1 and different concentrations of talazoparib for 5 days. Error bars are SEM for three independent experiments.

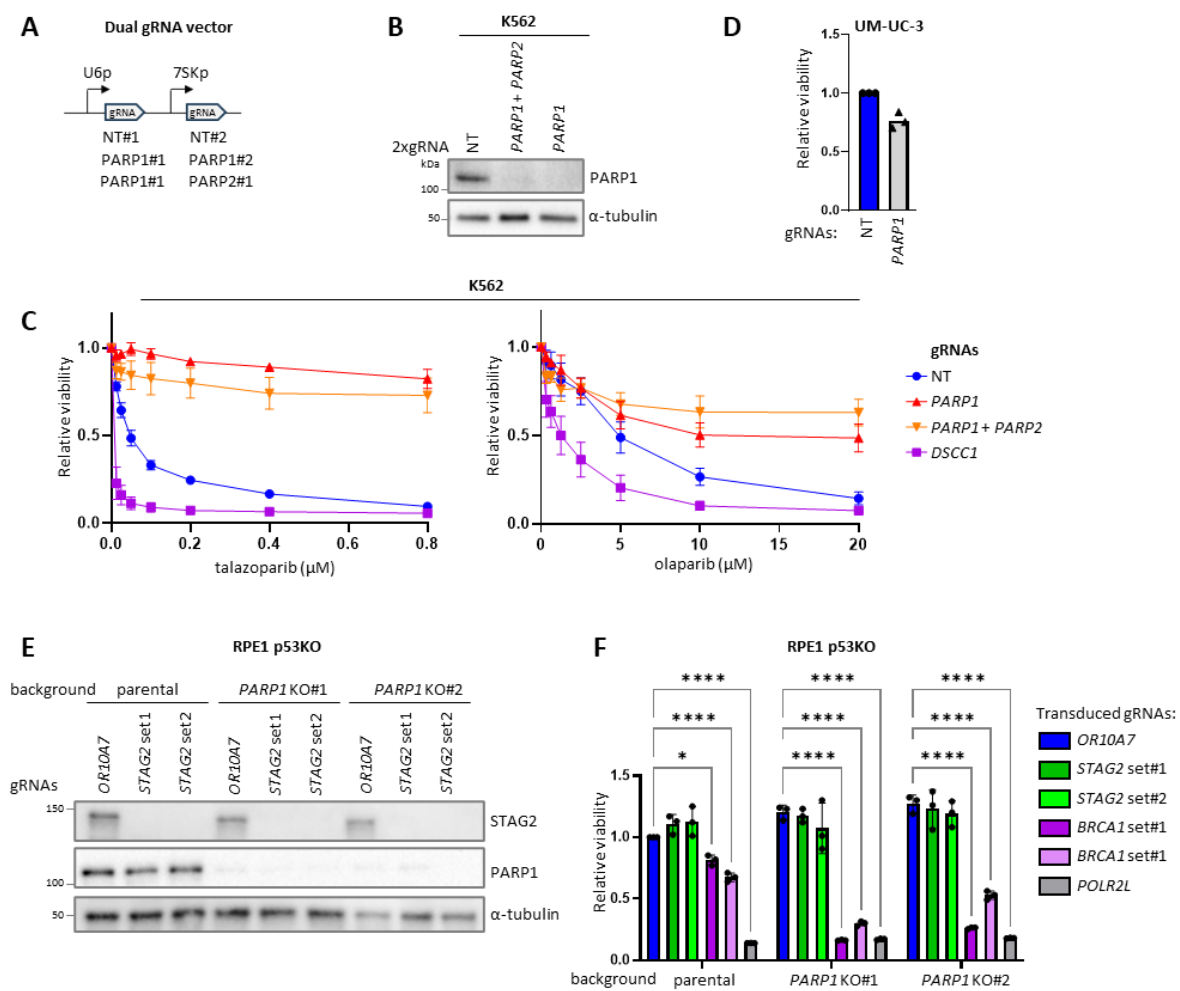

**Supplementary Figure 11. *PARP1* loss makes *STAG2*-deficient cells resistant to talazoparib and is synthetic lethal with *BRCA1* loss but not with *STAG2* loss.** (A) Schematic of dual gRNA expression vector for generation of (polyclonal) KO with Cas9. NT are a non-targeting gRNAs. (B) Western blot of K562 cells with *PARP1* or *PARP1* plus *PARP2* KO. (C) PARPi sensitivity of K562 *PARP1/2* KO cells. Cells were treated with PARPi for 6 days. (D) Relative viability of UM-UC-3 cells stably transduced with NT or *PARP1* gRNAs. Equal numbers of cells were seeded in a 96-well plate and viability was measured 5 days later. (E) Western blot of monoclonal RPE1 *PARP1* KO cells transduced with dual gRNA vector targeting *STAG2*. Two independent sets of gRNAs were used against *STAG2*. Guide RNAs against *OR10A7* are used as negative control. (F) Viability of *PARP1* KO cells transduced with gRNAs against *STAG2* or *BRCA1*. Transduced cells were selected with puromycin for 3 days and viability was measured 9 days after transduction. Error bars represent the SD of three independent experiments. Statistical significance was assessed using two-way ANOVA followed by Dunnett's multiple comparisons test.

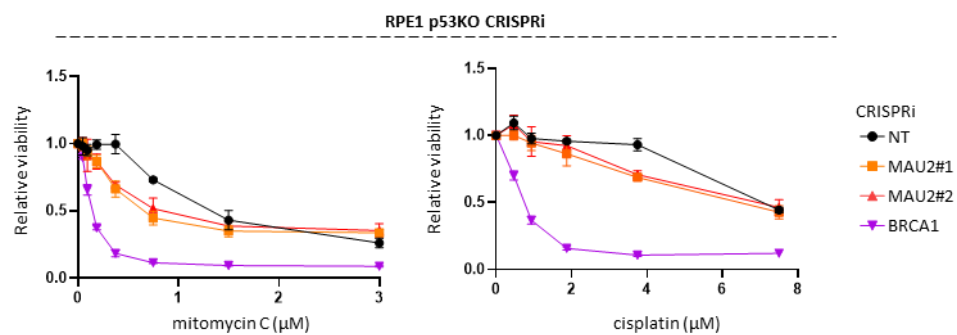

**Supplementary Figure 12. Sensitivity of *MAU2* KD cells to DNA interstrand cross-linkers.** RPE1 p53KO cells with CRISPRi against *MAU2* were treated for 5 days with cisplatin or MMC. Error bars are SEM for three experiments.

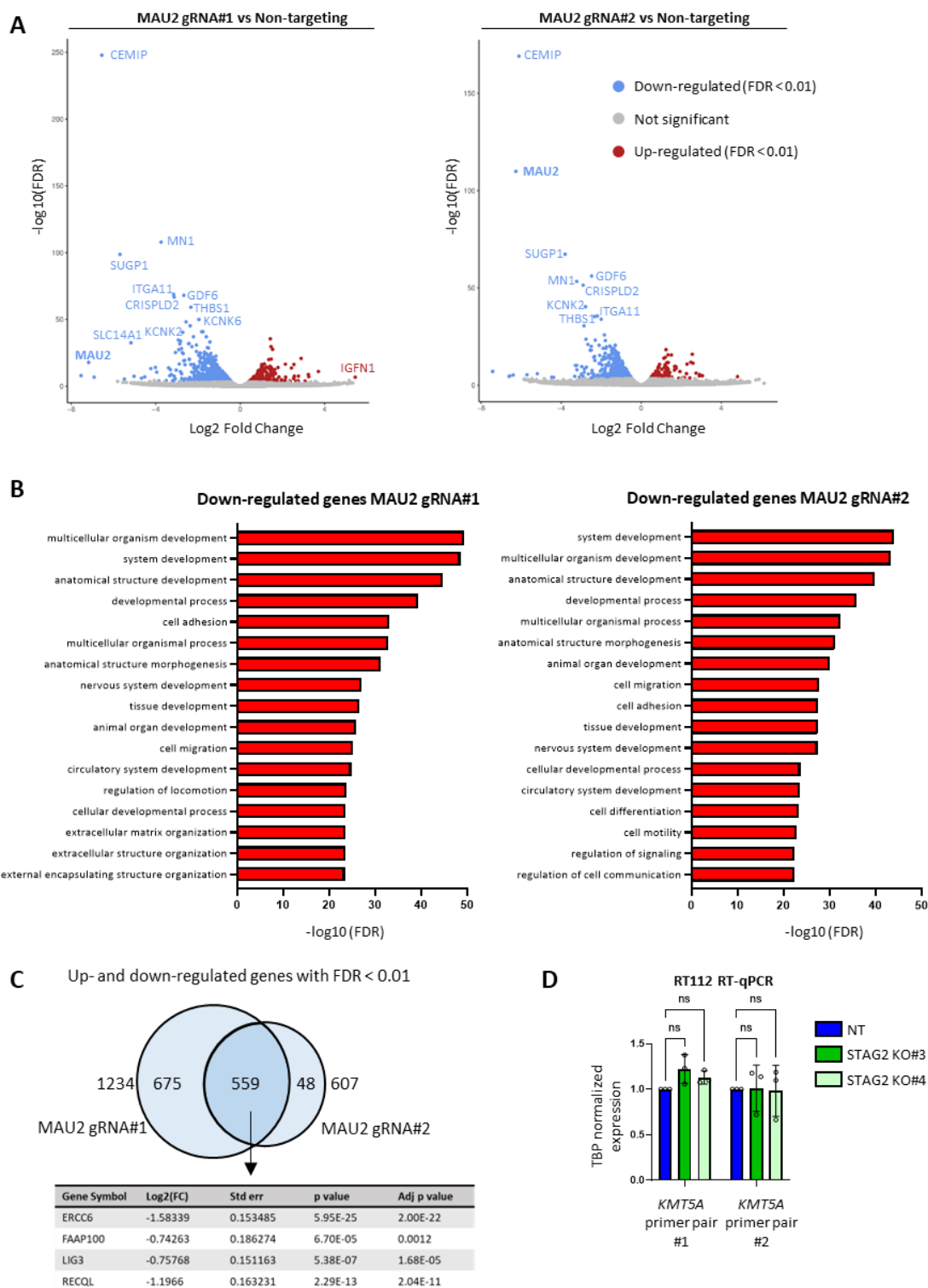

**Supplementary Figure 13. RNA-seq of MAU2 KD cells. (A)** Volcano plot of differentially expressed genes in RPE1 p53KO CRISPRi MAU2 KD cells versus cells with non-targeting gRNAs. **(B)** Gene ontology (GO) term enrichment of down-regulated genes. **(C)** Overlap of differentially expressed genes found in both sets of MAU2 gRNAs. *ERCC6*, *FAP100*, *LIG3*, and *RECQL* are DNA damage response related genes found in both gRNAs. **(D)** RT-qPCR for *KMT5A* in STAG2 KO RT112 cells. Each dot represents an independent experiment.

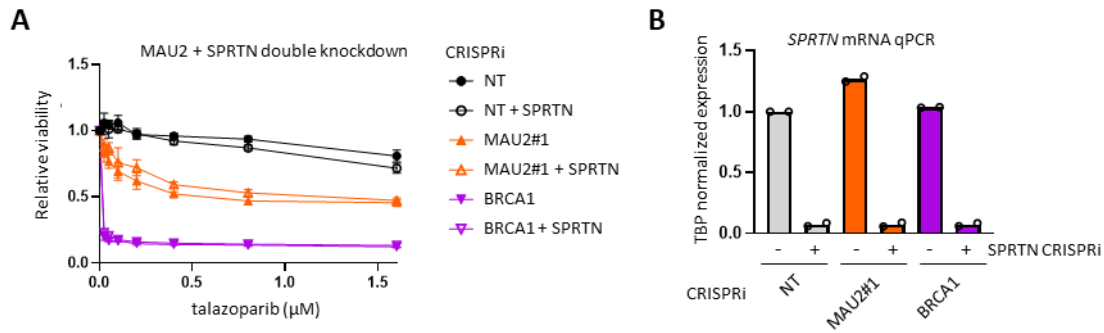

**Supplementary Figure 14. *SPRTN* KD does not further sensitize MAU2 depleted cells to talazoparib.**

**(A)** RPE1 p53KO cells with dox inducible CRISPRi against *MAU2* alone or *MAU2* and *SPRTN* simultaneously were treated with talazoparib for 5 days. Error bars represent SEM for three independent experiments. **(B)** RT-qPCR to confirm *SPRTN* downregulation after CRISPRi. Error bars represent the SD of three independent experiments. Statistical significance was assessed using two-way ANOVA followed by Dunnett's multiple comparisons test.

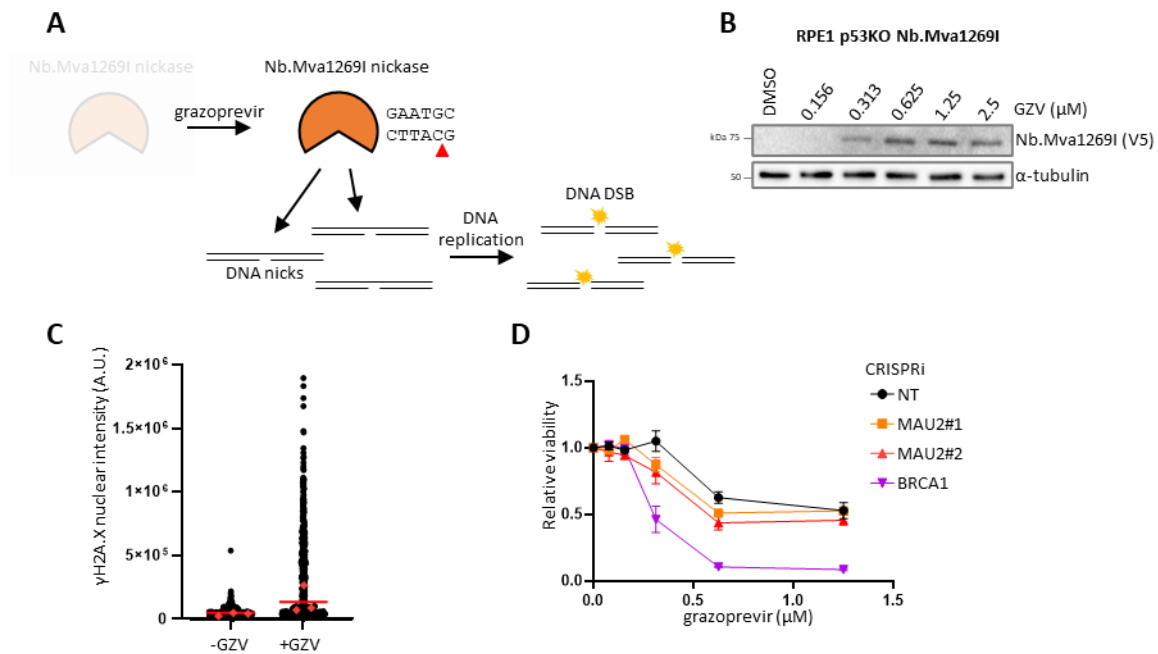

**Supplementary Figure 15. MAU2 KD cells show minor sensitivity to expression of the nickase restriction enzyme Nb.Mva1269I. (A)** Overview of GZV-inducible Nb.Mva1269I expression to generate DSBs in S-phase. The red arrow in the Nb.Mva1269I recognition sequence indicates the nick site. **(B)** Western blot of Nb.Mva1269I expression levels after 48h of treatment with different GZV concentrations. **(C)** Immunofluorescence of γH2A.X levels 72h after Nb.Mva1269I induction with 1 μM GZV. Each red dot represents the median γH2A.X intensity of at least 100 nuclei from one independent replicate, and the red bar represents the median of the independent replicates. **(D)** GZV sensitivity of CRISPRi cells with Nb.Mva1269I system. Cells were treated with GZV for 5 days. Error bars are SEM for three experiments.
